# Mutational signatures are markers of drug sensitivity of cancer cells

**DOI:** 10.1101/2021.05.19.444811

**Authors:** Jurica Levatić, Marina Salvadores, Francisco Fuster-Tormo, Fran Supek

## Abstract

Genomic analyses have revealed mutational signatures that are associated with DNA maintenance gone awry, a common occurrence in tumors. Because cancer therapeutics often target synthesis of DNA building blocks, DNA replication or DNA repair, we hypothesized that mutational signatures would make useful markers of drug sensitivity. We rigorously tested this hypothesis by a global analysis of various drug screening and genetic screening data sets, derived from cancer cell line panels. We introduce a novel computational method that detects mutational signatures in cell lines by stringently adjusting for the confounding germline mutational processes, which are difficult to remove when healthy samples from the same individuals are not available. This revealed many associations between diverse mutational signatures and drug activity in cancer cell lines, which are comparably or more numerous than associations with classical genetic features such as cancer driver mutations or copy number alterations. Validation across independent drug screening data and across genetic screens involving drug target genes revealed hundreds of robustly supported associations, which are provided as a resource for drug repurposing guided by mutational signature markers. We suggest that cancer cells bearing genomic signatures of deficiencies in certain DNA repair pathways may be vulnerable to particular types of therapeutics, such as epigenetic drugs.

## Introduction

Cancer precision medicine draws on the presence of somatically acquired changes in the tumor, which serve as predictive markers of response to drugs and other therapies. Commonly these markers are individual genetic changes, such as driver mutations affecting oncogenes or tumor suppressor genes, or copy-number alterations thereof. Many commonly employed cancer drugs act by interfering with DNA synthesis or maintenance or by damaging DNA. Therefore the altered capacity of cancer cells to repair and/or replicate DNA is the basis of many classical therapies, such as platinum-based agents, and also recently introduced or upcoming therapies, such as PARP inhibitors or ATR inhibitors. Therefore, it is paramount to derive predictive markers that are associated with failures of DNA maintenance in cancer cells. However, while DNA repair is often deficient in tumors, DNA repair genes such as MLH1 or BRCA1 or ATM do not commonly bear somatic mutations. Instead, they are commonly inactivated epigenetically, or by alterations in trans-acting factors, and so their deficiencies are difficult to predict from the genome. Additionally, germline cancer-predisposing variants may affect DNA repair genes, however pathogenicity of such variants is often challenging to predict. Because of the above, other types of molecular markers may be more useful to infer about failed DNA repair. This is exemplified in ‘BRCAness’ -- a gene expression signature that suggests a failed homologous recombination (HR) pathway, even in the absence of deleterious genetic variants in the BRCA1/2 genes.

In addition to gene expression, mutational signatures -- readouts of genome instability -- can characterize DNA repair deficiencies. One common type of signature describes relative frequencies of somatic single-nucleotide variants across different trinucleotide contexts. Certain mutational signatures were found to be associated with failures in DNA mismatch repair (MMR) and HR pathways^1^ as well as DNA polymerase proofreading^2,3^ and base excision repair (BER) ^4–6^ and nucleotide excision repair (NER)^7^ failures. Inducing DNA repair deficiencies in cancer cell lines is able to reproduce some of these signatures^8–11^. Other types of mutation signatures based on small insertions and deletions (indels)^12^ and on structural variants^13^ are also starting to be introduced.

Because mutational signatures describe the state of the DNA repair machinery of a cancer cell, they may be able to serve as a drug sensitivity marker. This was suggested recently for the “Signature 3” (also called SBS3), associated with pathogenic variants in BRCA1/2 genes^1,14^. SBS3 is common in ovarian and breast cancers, but it is also seen in other cancer types^15,16^, suggesting potential for broad use of drugs that target HR-deficient cells, such as PARP inhibitors. To this end, genomics-based predictors that draw on mutational signatures of HR deficiency have been developed^17,18^.

Human cancer cell line panels provide an experimental model for the diversity in tumor biology that is amenable to scaling up. Drug screens and genetic screens on panels of hundreds of cell lines^19,20^ proposed new cancer vulnerabilities by sifting through correlations between the sensitivity of cell lines to a drug (or genetic perturbation), and the genetic, epigenetic or transcriptomic markers in the cell lines. Encouragingly, genetic markers known to have clinical utility (e.g. BRAF mutations for vemurafenib, EGFR mutations for gefitinib, BCR-ABL fusion for imatinib sensitivity) are also evident in cell line panels^21^, suggesting their potential for discovery of useful genomic markers. Here, we investigated the hypothesis that mutational signatures evident in cancer genomes constitute markers of drug sensitivity using large-scale drug screening data sets of cancer cell lines.

However, quantifying somatic mutational signatures in cell line genomes is difficult, because a matched normal tissue from the same individual is typically not available and thus cannot be used to remove germline variation. A human cancer exome often contains two orders of magnitude more germline variants than somatic mutations (our estimates for several cancer types in Fig. 1A). This is typically addressed by filtering the known germline variants that occur in population genomic databases^22,23^. However, we estimate that after this, somatic mutations are still greatly outnumbered by the residual germline variants (Fig. 1B; a ∼9-fold excess, after removing all variants with population MAF>0.001%), which may confound downstream analyses such as the inference of mutational signatures. Here, we introduce a new method to infer somatic mutational spectra from cancer genomes without a matched control sample, while adjusting for the residual germline variation. We use this to infer trinucleotide mutation signatures in cancer cell line exomes, which are associated with sensitivity to drugs and to genetic perturbation across cell line panels. Replication analyses across independent data sets suggest that mutational signatures are remarkably broadly applicable markers of drug sensitivity, matching or exceeding common genomic markers such as oncogenic mutations or copy number alterations.

**Figure 1.**
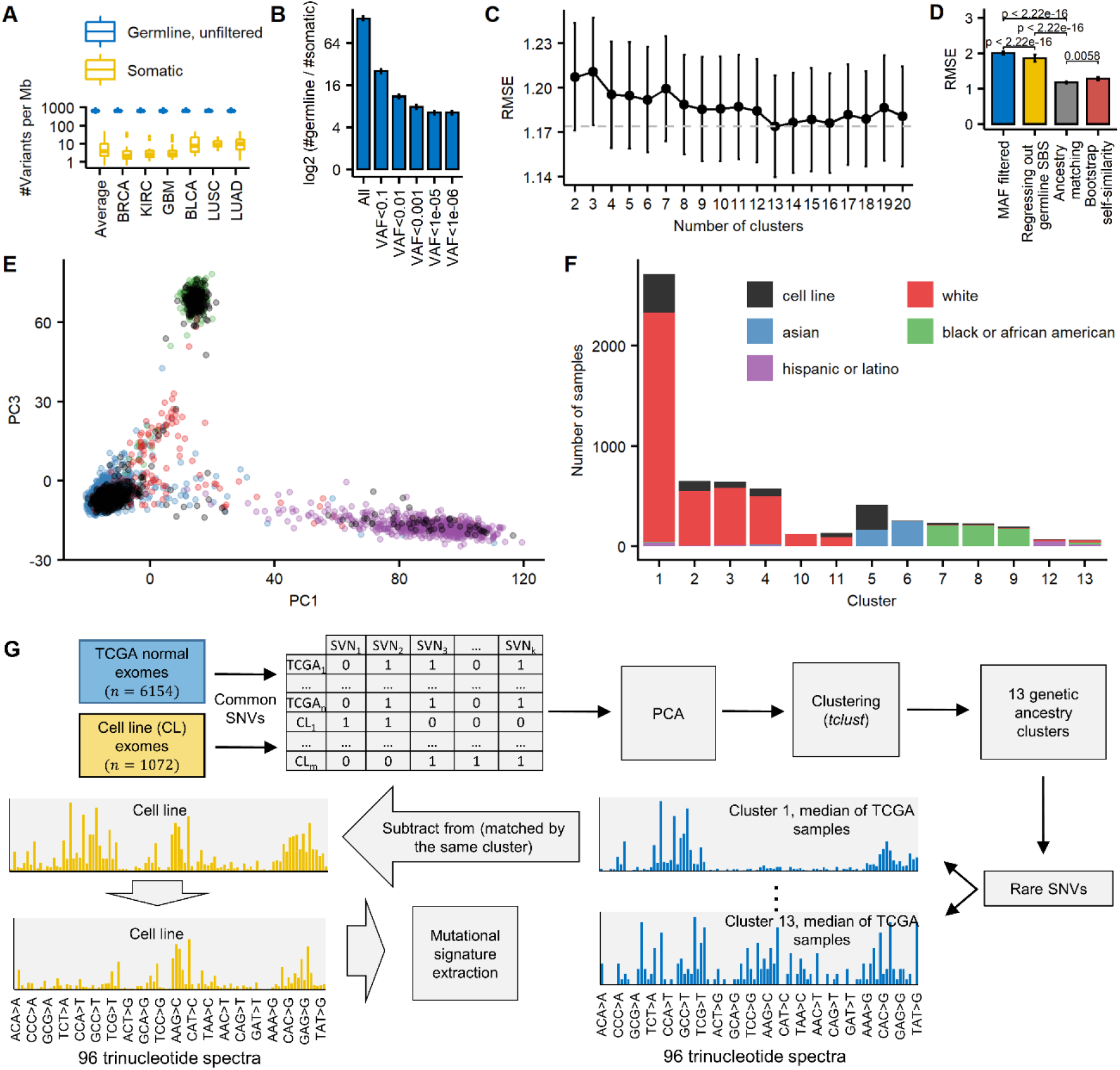
Evaluation of the ancestry-matching method to infer somatic mutation spectra on exomes without a matched normal control. (A, B) Germline variants greatly outnumber somatic mutations in exomes of various tumor types (A), after attempting to filter out germline variants according to the minor allele frequency (MAF) of variants found in the gnomAD database (B). (C) Root-mean-square error (RMSE) between the real somatic 96 tri-nucleotide profiles and the profiles obtained with the ancestry-matching procedure, after various number of clusters (based on principal components of common germline variants; see Methods) is considered. (D) Comparison of the ancestry-matching method (with number of clusters set to 13), the baseline procedure (variant filtering by population MAF<0.001%), regressing out mutational signatures reported as related to germline variants (signatures 1 and 5, and SNP signature^22,28^), and the RMSE expected by chance (estimated by bootstrapping mutations) P-values by Wilcoxon rank-sum test. (E) PC analysis of common germline SNVs (population MAF>5%) of TCGA samples and cancer cell lines recovers the major ethnicity groups. (F) Clustering based on the 140 main principal components (see Methods) reveals that cancer cell lines are represented across diverse genetic ancestry subgroups within the major ethnicity groups. (G) Schematic representation of the ‘ancestry matching’ procedure. Error bars in panels B, C, and D are the standard error of the mean.

## Results

### An ‘ancestry matching’ approach removes subpopulation-specific trinucleotide spectra to accurately infer mutation signatures

A substantial amount of the germline variation in a cell line exome cannot be removed by filtering based on minor variant frequency in population databases (Fig. 1B). Therefore we devised an approach to measure the somatic trinucleotide mutation spectrum -- the input for the inference of mutational signatures^24^ -- while rigorously adjusting for the contamination by the residual germline mutation spectrum.

Because germline mutational processes in human differ in different populations^25^, these differences have the potential to confound analyses of the somatic mutation spectrum. Since the residual germline variation in a filtered cell line genome can be very substantial (Fig. 1B), even slight differences in the germline spectrum can cause strong deviations in the observed spectrum, which is a mix of somatic and germline variation.

To address this challenge, we implemented an ‘ancestry-matching’ procedure, looking up, for each cell line genome, the individuals with a similar ancestry to the cell line’s ancestry. In particular, we clustered the cell line exomes together with germline exome samples from the TCGA data set, grouping by principal components derived from common germline variation (Figure 1G, Methods). The TCGA individuals provide a baseline mutational spectrum for that cell line, which can be subtracted from the observed spectrum to obtain the estimate of the somatic mutation spectrum.

We benchmarked our ancestry-matching procedure for accuracy of reconstructing the correct somatic mutation spectrum in a cancer cell line exome. To this end, we used single-nucleotide variant calls from TCGA cancer exomes where the matched normal was ignored to simulate the mutation calls that would be obtained from cell line genomes (see Methods). We then compared the reconstructed somatic mutation spectrum to the true somatic spectrum (obtained by contrasting tumor exomes with matched healthy tissue exomes from the same individuals).

Ancestry-matching greatly improves over the commonly used strategy, that is simply filtering out known germline variants according to population genomics databases (Fig. 1D, Supplementary Fig. 1D; root-mean-square-error (RMSE) is 1.174 *versus* 2.005; for comparison, the RMSE expected by ‘self-similarity’ with noise introduced via a bootstrap-resampling of mutations from the same samples, would be 1.277).

We considered various numbers of population clusters according to the RMSE in reconstruction of the correct somatic trinucleotide spectrum. Selecting three clusters, expectedly, recovers the major ethnicity groups (european, asian and african, Fig. 1E) and increasing the number of clusters to 13 clusters results in the minimum error in reconstructing trinucleotide mutation spectra (Fig. 1C; RMSE 1.174 for 13 *versus* 1.210 for 3 clusters).

Encouragingly, comparing the 13 ancestry clusters sorted by self-reported ethnicity (Fig. 1F), intra-ethnicity trinucleotide mutational profiles are more similar than the inter-ethnicity profiles (Supplementary Fig. 1A) and a PC analysis of the trinucleotide spectra of the rare variants separates the major ethnicity groups (Supplementary Fig. 1B). This is consistent with reports of differential mutagenic processes in the human germline across ancestral groups -- for example, the HCC>HTC (H = not G) variants were reported to be increased in Europeans, NCG>NTG mutations in Native Americans and NAC>NCN and TAT>TTT in some East Asians^25–27^. These reports, together with our benchmarking using simulations described above, support our rationale for using ancestry-specific baselines in inferring somatic mutational spectra of cancer cell lines (that do not have matched healthy samples available).

We applied the ‘ancestry matching’ methodology to exome sequencing data of 1071 cancer cell lines^29^, yielding their somatic trinucleotide spectra. On this data we performed *de novo* discovery using an NMF approach, broadly as described by Alexandrov *et al*.^24^ (with certain modifications, see Methods), where we extracted those NMF solutions that resembled previously reported tumor mutational signatures^12^ of single base substitutions (SBS). We have tested a number of variations on the data filtering and the mutation extraction methodology (Supplementary Table S1) to improve agreement with the set of SBS signatures^12^ as well to improve power of the set of mutational signatures to predict drug responses in the cell lines (Methods; Supplementary Table S1). We adopted a methodology that jointly infers trinucleotide (or SBS) signatures together with a set of indel features. Examining the SBS part of the spectrum, this yielded 29 cell line mutational signatures that very closely match (at a cosine similarity cutoff >=0.95) the known tumor SBS signatures, and further 21 cell line signatures that match known SBS tumor signatures (at a stringent cosine similarity >=0.85 and <0.95; a randomization test estimated that a >=0.85 cosine threshold corresponds to a 1.8% FDR in matching the correct SBS, Supplementary Fig. 4B).

The former group was labelled with the name of the corresponding SBS signature, while the latter similarly so plus the suffix “L” (for “like”). In some cases our cell line signatures were similar to more than one previous tumor SBS (Supplementary Fig. 4A) and they were named such as to make this evident, for instance our SBS15/6L matches two DNA mismatch repair (MMR) failure signatures (SBS15 and SBS6, more similar signature listed first). Note that a comparable degree of ambiguity is also observed among some of the known tumor SBS mutational signatures (Supplementary Fig. 5). The full set of 50 mutational signatures we inferred and their ‘exposures’ across cell types are visualized in Supplementary Fig. 2 and; Supplementary Fig. 3, and corresponding data is provided as Supplementary Tables S2 and S3.

Additionally, there were five mutational signatures that appeared specific to cell lines (SBS-CL), meaning they did not closely match one of the signatures from current tumor catalogues (Supplementary Fig. 2; Supplementary Table S2). These mutational processes may be evident only in rare tumor types or they may be active predominantly in cultured cells rather than in tumors. Some might originate from the incomplete separation of other signatures (Supplementary Figure 6 shows examples). Finally, some SBS-CL may reflect contamination with residual germline variation, as well as with sequencing artefacts, similarly as was recently reported for many SBS signatures recovered from tumor genomes^12^.

### Mutational signatures predict cell line drug response more accurately than oncogenic mutations or copy number alterations

Genetic and epigenetic alterations in cancer cell lines were on many occasions investigated as markers as sensitivity to chemical compounds^21^,^20^. We hypothesized that mutational signatures in a cell line genome can serve as similarly informative markers of drug sensitivity or resistance. We compared them to the predictive ability of the markers commonly used to predict drug response in cell lines: oncogenic mutations (in 470 cancer driver genes), recurrent focal copy number alterations (CNAs at 425 genes^21^), and DNA methylation data at informative CpG islands (HypMet at 378 genes^21^). Additionally we examined gene expression patterns (mRNA levels of 1564 genes that are either represented in the L1000 assay^30^ or are known drug target genes^31^), because gene expression was reported to be highly predictive of drug response^21,32^, possibly because it reflects the inherent differences in drug response between various cancer types and subtypes.

We predicted the sensitivity (log IC50 concentration) of a panel of 930 cell lines (separately for 29 cancer types that had a sufficient number of cell lines available) to a set of 518 drugs from the GDSC data set^29^. In particular we used Random Forest (RF) regression applied to the complete set of genetic or epigenetic markers in an individual cell line (listed above) as features (Figure 2A). In addition to mutational signatures inferred herein, we also considered the cell line mutational signatures reported by two recent studies^23^,^22^, obtained using approaches that did not account for ancestry and that have moreover fit the data to pre-existing sets of SBS signatures, rather than extracting new signatures from cell line data (see Methods).

**Figure 2.**
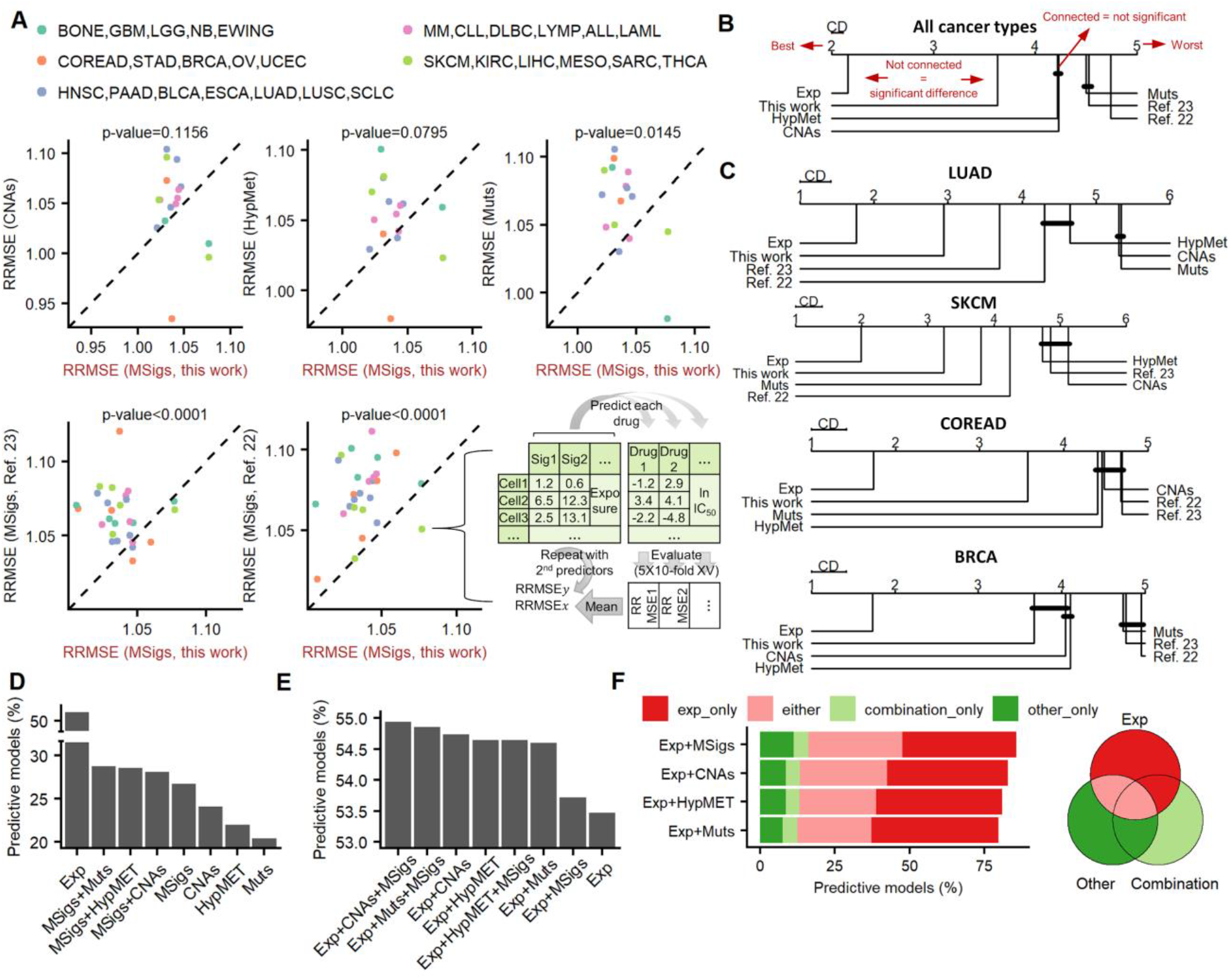
Prediction of drug response with mutational signatures and other molecular data types. (A) Predictive performance (RRMSE) of drug response prediction with mutational signatures (MSigs) reported here and previously^22,23^ and other data types (oncogenic mutations, copy number alterations and DNA hypermethylation). P-values of paired one-sided Wilcoxon signed rank test are also reported on the plots. Dashed line denotes the diagonal. (B) Average ranks diagrams for the performance (RRMSE) of the drug response prediction with various predictors. Each graph represents two different kinds of information: the ranking among the predictors (the predictors positioned at the leftmost side of each graph are the best performing) and the statistical significance of the difference between pairs of predictors: if the ranks of two predictors are at least critical distance (CD) apart, the difference in their performance is considered statistically significant (at p<0.05) according to the Nemenyi post-hoc test. The groups of predictors for which this does not hold are connected by black lines. (C) Tests shown separately for four tissues-of-origin with the highest number of cell lines in our panels. (D) The percentage of the Random Forest models that are predictive of drug response, defined as having a predictive error lower than the one of an uninformative default model (predicts the average log IC50 for every cell line). Expressed relative to the total number of testable drug-tissue pairs. (E) Percentage of the predictive models obtained by combining gene expression with other features. (F) The percentage of models that are predictive with gene expression but not another feature type (“exp_only”), by other feature type but not with gene expression (“other_only”), by either model (“either”), or only by a combination of both (“combination”).

Firstly, mutational signatures predicted drug sensitivity significantly better than all other tested types of alterations: increase in accuracy of RF models over CNAs, DNA hypermethylation, and oncogenic mutation features is significant (at p<0.05, by corrected Friedman test followed by the post-hoc Nemenyi test on ranks; Fig. 2B; Methods; average rank for mutational signatures was 3.63 while for other types of (epi)genetic features it was 4.22–4.50 considering all RF models). Secondly, mutational signatures found by our ‘ancestry matching’ approach perform significantly better than other cell line signatures recently reported^22,23^, comparing on the set of cell lines that overlap between the publications (Fig. 2B, C). Moreover, these previous sets of cell line mutational signature exposures were less predictive of drug sensitivity than were CNAs, oncogenic mutations and DNA methylation, suggesting the utility of adjustment for germline spectra contamination prior to mutational signature inference (Fig. 2B, C, D).

Next, we applied a different test that considers the average error in predicting the drug sensitivity profile (relative RMSE of a RF model in crossvalidation; Fig. 2A), averaged across all drugs in a given tissue. In most cancer types, the mutational signatures obtained herein were better predictors of drug response, than all the usual genomic and epigenomic features (11 out of 16 tested cancer types comparing with DNA hypermethylation, 13 out of 16 for CNAs, and 12 out of 16 for oncogenic mutations). Also, in most cancer types (Fig. 2A) our cell line signatures significantly outperformed recent methods to infer mutational signatures^22,23,33^ that are naive to germline mutational spectra (in 21 and 26 out of 27 cancer types for the two previous methods that used the same set of cell lines, Fig. 2A, and in 11 out of 16 cancer types on a set of overlapping cell lines for a third method (Supplementary Fig. 7A); p<0.0001, p<0.0001, and p=0.037, respectively, Wilcoxon test for decrease of relative RMSE).

Gene expression was overall very highly predictive of drug response, (Fig. 2B, C, D), consistent with recent reports^21,32^. We asked if the predictive power of gene expression can be complemented by additionally including mutational signatures and/or various sets of genetic markers. We predicted the drug response profile in RF models as above (Fig. 2A), but here by using combinations of marker types with gene expression, and tallying the predictive RF models (drug-tissue pairs with better-than-baseline RMSE in crossvalidation; Methods). Notably, gene expression is complemented by mutational signatures and also by other types of features, yielding a higher percentage of predictive RF models when markers are combined than with gene expression alone (Fig. 2D, E), with the optimal combination being gene expression with mutational signatures and with CNAs (Fig. 2E). Even if gene expression markers are unavailable, the mutational signatures were still complementary to oncogenic mutations, CNAs, or DNA methylation (Fig. 2D).

Next, we considered the complementarity analyses at the level of individual drugs, asking if the profile of drug sensitivity that can be predicted by a combined RF model (e.g. gene expression and mutational signatures) could also have been predicted by the two RF models drawing on the individual sets of features – on gene expression only, or on mutational signatures only (Fig. 2F). The number of RF models where gene expression by itself is not predictive but mutational signatures are predictive is substantial (1265 drug-tissue pairs, plus 566 where only a combination of signatures and expression is predictive). This is higher than the number of drug-tissue pairs where gene expression is not predictive but driver mutations are (563 plus 352), and similarly so for CNA (633 plus 347). In addition to gene expression, using DNA methylation as a baseline also supports that response profiles for many drug-tissue combinations can be predicted only by mutational signatures (Supplementary Fig. 7B). This suggests that the predictive signal in mutational signatures does not simply reflect cancer subtype or cell-of-origin, at least to the extent that subtype can be identified via gene expression or DNA methylation patterns. Overall, mutational signatures, considered collectively, can complement gene expression and other types of markers in predicting drug sensitivity profiles of cancer cells.

### Many associations between individual mutational signatures and response to drugs

The above analysis (Fig. 2) indicates that SBS mutational signatures considered in aggregate yield an accurate predictor of differential drug response across cancer cell lines. However it is interesting to establish associations between individual mutational signatures and response to particular drugs to be able to apply mutational signatures as genomic markers. Additionally such associations might suggest the underlying mechanism of cell killing or growth inhibition by the drug, based on the known aetiology of a particular signature.

Therefore we have performed statistical tests for associations, using a previous approach based on ANOVA^21^ to link drug sensitivity with different cancer functional events (CFE, here implying CNAs, HypMet and oncogenic mutations), herein applied to link drug sensitivity and mutation signatures (a continuous value of the relative NMF exposure i.e. normalized to the sum of exposures of all signatures in that cell line; Supplementary Table S3; Methods). We tallied the associations with FDR<=25% in every cancer type separately. Across the 518 drugs, we found 502 associations (Supplementary Table S5), while for oncogenic mutations we found 154, for copy number alterations 160, and for DNA methylation 74 significant associations (Supplementary Table S5; we note that the number of testable associations with mutational signatures was 1.7-fold higher than for the other markers because of lower sparsity of the marker values for signatures; we required at least 3 non-zero values per tissue). This suggests that mutation signatures, considered individually, can provide highly informative genomic markers of sensitivity or resistance to many drugs.

The significant associations encompassed all 55 mutational signatures: each had at least one drug association at FDR<25%, including all five SBS-CL (Supplementary Fig. 8). This further supports that SBS-CL are indeed *bona fide* mutational processes not artefacts. Moreover, the significant associations with mutational signatures were detected across 21 different cancer types (Supplementary Fig. 8). The ability of the mutation signatures to predict a drug sensitivity profile was rarely explained by the total mutation burden in the cell ine (Supplementary Fig. 7B).

### Associations with drug response that replicate in independent data sets

Because of reproducibility concerns in large-scale drug screens^34–36^ that might stem from technical reasons or from cancer cell line evolution during culture, we asked if the associations between mutation signatures and drug responses replicate across data sets. To this end, we tested for associations involving mutational signatures or other markers, with the additional condition that they also replicate in an independent data set. We implemented a randomization-based procedure that tests that the smallest effect size (Cohen’s *d* statistic) across both datasets is above chance (Fig. 3A). A value of *d*>=1, typically considered a large effect size, implies that a difference in mean drug sensitivity (log IC50) between the cell lines positive for a feature and those negative for a feature is greater than the pooled standard deviation (of the log IC50) of the two sets of cell lines. These replication tests were performed on binarized mutational signatures i.e. the signature present/absent indicator variables, considering different cancer types individually (Supplementary Table S4). Additionally the same tests were performed on the usual markers in cell line screening analyses, including oncogenic driver mutations, CNAs and promoter DNA methylation.

**Figure 3.**
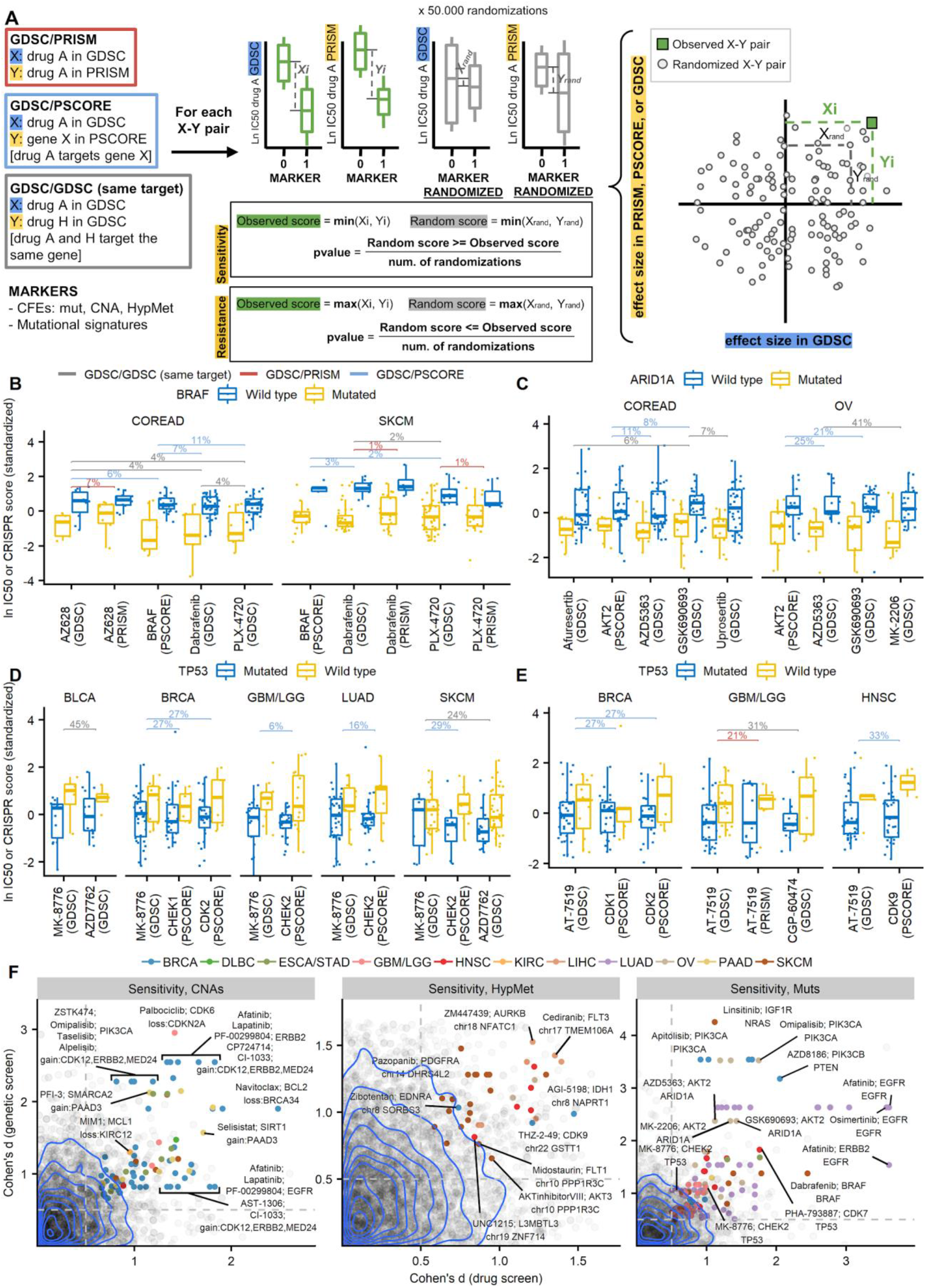
Detecting robust associations between genetic or epigenetic markers and drug sensitivity by replication across measurements. (**A**) The randomization test methodology to detect replicating associations using three different tests: consistent effects of a drug between two screening assays (GDSC/PRISM), effects of a drug consistent with effects of knockout of the target gene (GDSC/PSCORE), and effects consistent across different drugs that share the same molecular target (GDSC/GDSC or “same target”). (**B**-**E**) Examples of replicated associations of a known example of oncogene addiction (to *BRAF*, **B**) and of novel examples of cancer vulnerabilities associated with mutations in tumor suppressor genes (*ARID1A*, **C**; *TP53* **D** and **E**). Y-axes show a Z-score derived from either the ln IC50 value (drug sensitivity) or from the CRISPR essentiality score, depending on column. Horizontal bars show false discovery rates (q-value adjusted) obtained via a randomization test in panel A, where color denotes the type of the test. (**F**) Effect sizes of markers that associate with drug response in the GDSC drug screen (X-axes) and with response drug target gene knock-out in the Project SCORE genetic screen (Y-axes), shown separately for copy number alterations (CNA), promoter DNA hypermethylation (HypMet) and mutations in cancer genes (Muts). Gray points represent all tested associations, while colored points denote the statistically significant associations that also meet an effect size threshold. Blue lines are the contours of the 2D kernel density estimates. Representative points are labeled. Drug names on grouped labels (the CNA sub-panel) are ordered by their appearance on the plot from left to right.

We considered three different types of replication methods: an external replication in an independent drug screening data set, internal replication with multiple drugs affecting the same target, and a replication using CRISPR/Cas9 gene knockout fitness data. Randomization p-values from these replication methods across various tissues were rarely inflated for mutational signatures, and in fact commonly exhibited slight deflation (mean lambda across tissues 0.57– 0.95 for different replication methods, Supplementary Fig. 9) suggesting an overall conservative bias in the replication test as implemented.

Firstly, we performed a replication analysis where the drug association data from the GDSC was tested against another drug screening data set: PRISM (derived from a different experimental methodology involving pooled screens^37^; 348 cell lines and 178 drugs overlap with the GDSC set). In total 223 drug-mutation signature associations were robustly supported across both GDSC and PRISM (*d*>=0.5 and same direction of effect in both datasets and additionally requiring randomization test FDR<25%; adjustment using the q-value method^38^), observed across diverse tissues and diverse signatures (Figure 4A, B; Supplementary Figure 10A). This matches or exceeds the number of drug associations replicated in PRISM involving driver mutations (86) and copy-number changes (178), DNA methylation (261) in the same test. We list all replicated associations in Supplementary Table S6.

**Figure 4.**
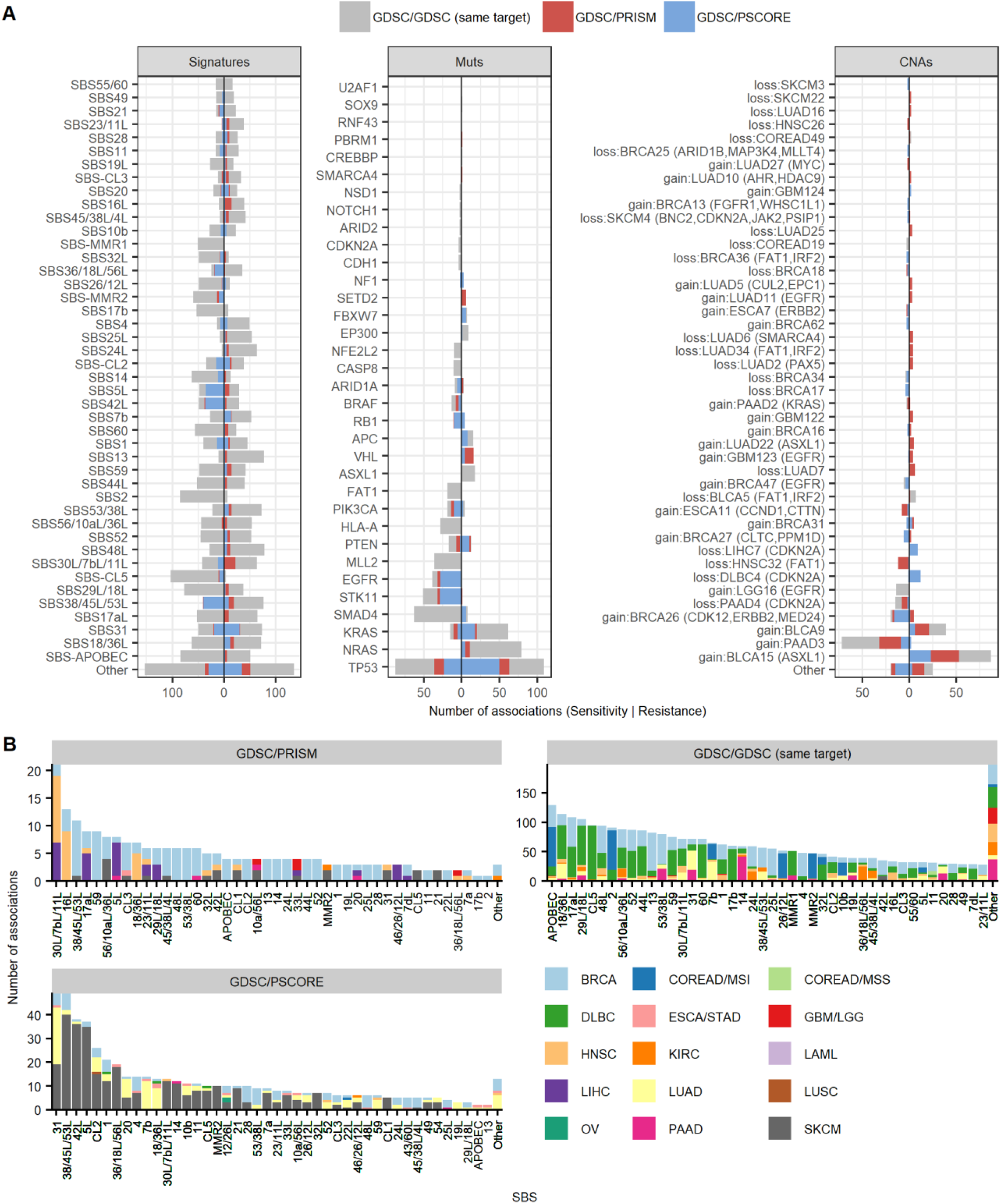
Tally of the significantly replicated associations of drug sensitivity or resistance with mutation signatures and other markers. (A) Comparison of the number of statistically significant associations (FDR<25% by randomization test, additionally requiring an effect size *d*>0.5) per feature, among mutational signatures, oncogenic mutations and copy number alterations in the three types of replication tests (see Fig. 3A). Features are ranked by the number total of significant associations, either for drug sensitivity (negative side of X-axis) or resistance (positive side of X-axis). (B) Distribution of statistically significant associations across different cancer types and binarized mutational signatures in the three replication tests.

Given that the amount of cell lines available to the replication analysis is smaller and thus statistical power is limiting particularly for some tissues, we suggest that some associations at permissive thresholds (here, nominal p<0.01) may be of interest. This is illustrated by a known example where MMR failures associate with sensitivity to DNA topoisomerase I inhibitors, based on prior experimental work in colon cell lines^39^ and on clinical data from colon cancer patients^40^. In our analysis, the MMR signature SBS26/12L predicted sensitivity to camptothecin in colorectal cells, which replicated in PRISM at p=0.009 (at FDR=23%, Supplementary Table S7). Additionally, we found SBS20-associated topotecan and SN-38 sensitivity in skin (topotecan sensitivity also replicates for SBS21), and SBS14 SN-38 sensitivity in pancreas. All these replicated in PRISM at p=0.0067 or lower but at FDR>25% thus supporting the careful usage of our replication analyses also at the permissive cutoff (note that further replicated links between MMR signatures and sensitivity to topoisomerase I drugs were found in additional tissues such as lung; list in the Supplementary table S7).

Secondly, we performed an internal replication analysis within GDSC, enforcing that associations must be detected with two or more drugs that share the same molecular target. In total, 228 drugs in the GDSC data could be tested in this ‘same-target’ analysis. Effectively, multiple drugs serve as pseudoreplicates, and additionally this test helps discard associations due to off-target effects, which are more likely to differ between two drugs than their on-target effects. Here, we found 2849 significant associations for mutation signatures, 661 for oncogenic mutations, 171 for copy-number changes, and 970 for DNA methylation (at effect size d>0.5 and FDR < 25%) (Figure 4A, B; data in Supplementary Tables S8 and S7 for the associations between the default FDR<25% threshold, and the permissive threshold at p<0.01, respectively).

### Robust associations that replicate multiple types in different tissues or testing methods

Some associations overlapped between the same-target and the GDSC-PRISM replication analyses, suggesting more robust associations: Supplementary Table S7 contains those replicated associations that were seen either across multiple cancer types, and/or across multiple drugs that target the same pathway, and/or across different replication methods (including also CRISPR genetic screens, see below). We suggest that this ‘silver set’ of 13846 associations, where each replication at a FDR<=25% is supported by one or more additional replications at a suggestive (p<0.01) threshold, to be potentially suitable for further analyses or follow-up.

For example, mutations in the MLL2/KMT2D gene in colorectal cancer cells, which associate with sensitivity to the Aurora kinase inhibitors GSK1070916, ZM447439, CD532 in GDSC data (FDR=4%-6%). The GSK1070916 is also replicated in PRISM in colorectal cells (FDR=13%), Supplementary Table S7). This suggests that the loss-of-function of the KMT2D chromatin modifier may generate vulnerabilities to Aurora kinase inhibition in diverse tissues, which mirrors recent experimental data in cell line models^41^. Mutations in the MLL2/KMT2D gene also sensitize to the bromodomain inhibitors OTX015 and I-BET-762 (FDR=10%) as well as PFI-1 and RVX-208 (FDR=10%) in GDSC data. The OTX015 association with MLL2 mutations also replicates in PRISM data (FDR=16%). In another cancer type (bladder cancer), MLL2 mutations sensitize to both OTX015 and JQ1 bromodomain inhibitors at a permissive threshold (p=0.0033).

Intriguingly, a large number of significant associations in the same-target replication test involved resistance to diverse drugs in NRAS-mutant acute myeloid leukemia cells (Supplementary Table 8). NRAS mutations in myeloid leukemia predispose to resistance to many different drugs (GDSC data^21^ shown in Supplementary Figure 11) and have thus resulted in many non-specific hits in our replication test (n=89, Figure 4A). This result is consistent with the reported association of NRAS mutant blood cancers with poor prognosis, drug resistance and relapse^42–44^ suggesting that NRAS mutations indeed confer a generalized drug resistance phenotype in myeloid cells.

### Integrating drug screening data with genetic screening data to obtain robust associations

As a third type of replication analysis, we further prioritized cancer vulnerabilities by intersecting the drug sensitivity screening data sets with genetic screening data sets across a panel of cell lines. Our rationale was that a biological process may be targeted similarly by pharmacological inhibition of a protein that performs the process, or by editing the genes that encode the corresponding protein. In specific, we examined sensitivity to CRISPR/Cas9-mediated knockouts that target the protein-coding gene across a panel of 517 cell lines^45^ that overlapped the GDSC cell lines. We discovered many associations with conventional markers -- oncogenic mutations (n=243), copy number alterations (n=148), and DNA methylation (n=100) -- that replicated across drug and genetic data (Supplementary Table S9, Figure 4, Figure 5A, Supplementary Figure 10B).

**Figure 5.**
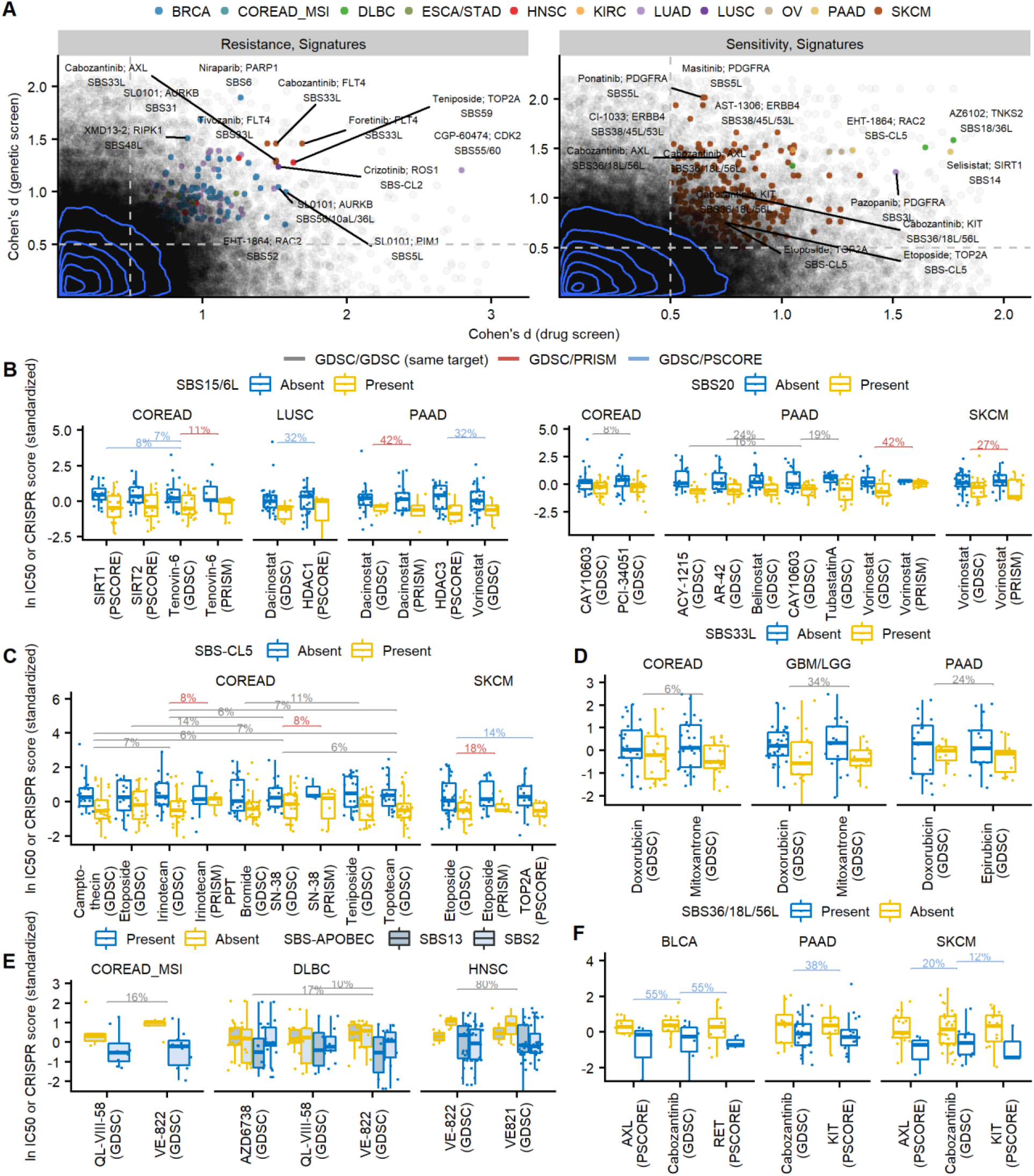
Associations with drug sensitivity or resistance that replicate in independent datasets. (A) The GDSC/PSCORE replication test of mutational signatures that robustly associate with drug response (X-axes) and also with response to the knock-out of the drug target gene (Y-axes). Gray points represent all tested associations, while colored points denote the statistically significant associations that also meet the effect size threshold of *d*>0.5. Blue lines are the contours of the 2D kernel density estimates. Representative points are labeled with drug name, gene name and SBS mutational signature involved in the association. See Fig. 3F for related plots for other types of markers. (B-F Examples of associations of mutational signatures with drug sensitivity that replicated (using tests in Fig. 3A) multiple times, across different cancer types and/or different types of replication tests. Y-axes show a Z-score derived from either the ln IC50 value (drug sensitivity) or from the CRISPR essentiality score, depending on column. Horizontal bars show false discovery rates (q-value adjusted) obtained via a randomization test in panel A, where color denotes the type of the test.

Demonstrating the utility of this test, we recovered known examples of the oncogene addiction paradigm, such as the breast cancer and esophageal/gastric cancer cell lines with ERBB2 (HER2) amplification being sensitive to inhibitors of EGFR and ERBB2 (lapatinib and CP724714), and also to the ERBB2 gene knockout (replication FDRs all <=21%). Similarly, the BRAF mutations in skin and colorectal cancer likewise sensitize to different BRAF inhibitors (e.g., dabrafenib, AZ628, PLX-4720 in both the GDSC and PRISM) and also BRAF gene disruption (Fig. 3B); this and other striking associations with gene mutations, CNA and promoter DNA methylation that replicate are highlighted in the global data overview in Fig. 3F.

In addition to oncogene addiction, replicated associations may suggest ways to target mutated tumor suppressor genes *via* synthetic lethality. One example involving a known pathway is that the CDKN2A deletion sensitizes glioblastoma/glioma cells to palbociclib (a CDK4 and CDK6 inhibitor), and this is also recapitulated in the knockout of the CDK6 gene (FDR=14%, p=0.00006) and also suggestively with the knockout of CDK4 (at FDR>25%, p=0.0005). CDKN2A loss as a biomarker for intervention on CDK4/6 was proposed by previous work on preclinical models^46–48^ and is consistent with the biological role of the CDKN2A-encoded protein p16^INK4a^ in inhibiting CDK4/6. In contrast to CDKN2A, mutations in the tumor suppressor RB1 were associated with insensitivity to palbociclib and also to CDK4 and 6 gene knockout in glioma/glioblastoma (FDR=6% and 1%, respectively). This is consistent with the role of CDK4/6 in inactivating RB1 to promote S-phase entry and with reports that RB1 mutation predicts resistance to CDK inhibitors^47,49^. Overall, this synthetic lethality example solidifies previous findings on targeting *CDKN2A* or *RB1* mutant tumors by CDK inhibition and demonstrates the potential of the joint analyses of drug and genetic screening data to uncover cancer vulnerabilities, suggesting that other associations we recovered (Supplementary Table S9) might provide promising avenues for treatment.

We highlight an unanticipated and robustly supported synthetic lethality example involving mutations in the *ARID1A* tumor suppressor gene and the inhibition of the AKT2 gene or protein (Fig. 3C). In particular, *ARID1A* mutant colorectal cell lines are more sensitive to the knock-out of the *AKT2* gene by CRISPR, as well as to the pan-AKT inhibitors, GSK690693 and capivasertib/AZD5363 (FDR=8% and 11% in the replication test, respectively). The same is observed in ovarian cancer cell lines, again involving *AKT2* knockout and the same two inhibitors (at FDR=20% and 25%, respectively). This is supported by additional AKT inhibitor drugs: afuresertib (FDR=11%/p=0.0002), AKT inhibitor VIII (FDR=20%/p=0.0051), and uprosertib (FDR=11%/p=0.0002) in colon, and MK-2206 (FDR=21%/p=0.0013) in ovary (Supplementary Table S9). The *AKT2* oncogene can be amplified in ovarian, endometrial, pancreatic and other cancer types, while the *ARID1* tumor suppressor commonly bears truncating mutations in many cancers. In tumor genomes, *AKT2* alterations significantly co-occur with *ARID1A* alterations (OR=2.0, FDR<0.1% in MSK-IMPACT cohort of 10,945 samples^50^; replicated at OR=1.4, FDR<0.1% in TCGA pan-cancer cohort of 10,967 samples; analysis via cBioPortal^51^), suggesting that the *AKT2* amplifications may bring a selective benefit to *ARID1A*-mutant cancers. Our analyses provide robust support to the notion that the PI3K/AKT/MTOR signaling inhibition may be a vulnerability of *ARID1A*-mutant tumors^52–55^ and further suggest specifically AKT2 as an opportune point of intervention.

Further, our analyses suggest a robust association of *TP53* mutation status with sensitivity to CHK and CDK kinase inhibitors across four different cancer types (at a FDR<30%, and two additional cancer types at a permissive threshold; Fig. 3D, E). MK-8776 is an inhibitor of the CHK1 protein, with some activity against CHK2, which we found differentially active against TP53-mutant cells. In the majority of cancer types however the TP53 mutation sensitized to knockout of the secondary target *CHEK2* gene (Fig. 3D; we did observe *CHEK1* in breast). This suggests that the off-target activity of MK-8776 towards CHK2 contributes to the killing of TP53 mutant cells, further supported in the activity of the AZD7762 drug (in breast cell lines, Fig. 3D) which inhibits CHK1 and CHK2 to similar extent. In a similar vein, AT-7519, a multi-CDK inhibitor with preferential activity towards CDK9 and CDK5, is more active upon *TP53*-mutant cell lines of three tissues (Fig. 3E). However, our analysis suggests a key role for AT-7159 activity towards CDK2, evidenced via sensitivity to CDK2 k.o. in breast, and to the CGP-60474, an inhibitor more potent against CDK2, in brain cells (Fig. 3E). These examples suggest vulnerabilities of *TP53*-mutant cells to manipulating the activity of CHK2/CDC25A/CDK2 axis, as well as highlighting examples of how gene editing can reveal that drug effects are often exerted via secondary or unintended targets^56,57^.

Next, we applied this same statistical methodology (Fig. 3A) to analysis of mutational signatures in cell line genomes.

### Mutational signatures concurrently associated with sensitivity to pharmacological and to genetic perturbations

We studied the capacity of mutational signatures to predict fitness changes upon genetic perturbation. As a positive control, we expected to recapitulate a recently reported vulnerability of cell lines that are microsatellite-instable (MSI), and therefore deficient in the DNA mismatch repair (MMR) pathway, which do not tolerate a loss of the *WRN* gene^45,58–60^. MMR deficiencies in tumor genomes associate with MSI, and also with trinucleotide mutational signatures 6, 15, 21, 26 and 44^12^ (and additionally 14 and 20 which result from MMR failure concurrent with deficiencies in replicative DNA polymerases). We found links between many of the corresponding SBS that we inferred in the cell line exomes and sensitivity to *WRN* knockout in a joint analysis of MSI-prone cancer types (colorectal, ovary, stomach, uterus), however, with different levels of support (FDRs 29%, 90%, 98%, 1.2%, <0.01%, 29%, 13% and N.S. [associated with resistance]) for SBS6, 12/26, 14, 15/6L, 20, 21, 26/12L and 44L, respectively; Supplementary Fig. 9A). Additionally, two signatures consisting of A>G changes that also bore a high weight on the indel component and thus might be MMR-related, SBS33L and SBS54 (Supplementary Fig. 2), also predicted sensitivity to *WRN* loss (Supplementary Figure 12). Our data thus suggests that some MMR-associated signatures (particularly SBS15, 20 and 26) are more robust markers for WRN inhibition than the other MMR failure-associated signatures (such as 6, 14, 21 or 44), possibly because the SBS reflect different modes of failure of the MMR machinery that confer differential requirements for WRN. Beyond *WRN* loss, the MMR and other mutation signatures can also predict sensitivity to many drugs, including some that the MSI status would not predict (Supplementary Fig. 7B; we note that the converse is also true, at least for the few cancer types where MSI labels are available).

Recent work suggested that MMR-failure SBS signatures can be grouped into a few broad types^61–63^, suggesting a signature group enriched with C>T changes, and a group enriched with A>G changes. Based on this we further tested MMR signatures for drug associations both individually and also as a part of two groups (MMR1 is a sum of the SBS6, SBS15/6L and SBS44L exposures, while MMR2 is a sum of the SBS26/12 and SBS21 exposures; Supplementary Figure 12). Overall, the ability to recover the known *WRN* dependencies of MMR-deficient cell lines -- here, estimated via trinucleotide mutational signatures -- supports the utility of our methodology to infer mutational signatures in cell lines.

We systematically examined all mutational signatures for the overlap between associations in the GDSC drug screen and Project SCORE genetic screen. This yielded 540 associations (at a randomization FDR<25%, and additionally requiring an effect size of *d*>0.5 in both the genetic and the drug screens) that replicated across data sets -- a higher number than for oncogenic mutations, CNAs, and DNA methylation (243, 148, and 100, respectively). The associations involved 52 different signatures (Figure 4, Figure 5A; full list in Supplementary Table S9) and 114 genes. This broad prevalence of mutational signatures that predict drug and genetic screen responses is similar to the replication analyses described above, which considered either different drugs with the same target, or when considering replication across independent screens. The number of replicated associations involving highly-ranked mutational signatures by the Project SCORE replication analysis (such as the oxidative damage repair signature SBS18/36L, the chemotherapy pretreatment-associated signature SBS31 and the common signature of unknown aetiology SBS5L) Figure 4B, Supplementary Figure 10A, n=13, 49 and 37 replicated associations, respectively) broadly matches the replicated associations involving common oncogene mutations such as EGFR or BRAF or PIK3CA (n=28, 4, and 12, respectively), or known copy number change driver events such as ERBB2 gain (n=17 replicated associations in Project SCORE) (Figure 4A; Supplementary Table S9). We note that in this and further analyses we have, conservatively, stratified colorectal cancer cell lines into MSI and MSS (association counts without stratification are in Supplementary Fig. 13; Supplementary Table 10). This is because MSI is common in colorectal cell lines, and the MSI status is expected to be strongly associated with MMR signatures^12,63^ and possibly other mutational signatures^64^.

### Mutational signatures of DNA repair failures predict response to epigenetic drugs

We examined the list of replicated drug associations involving mutational signatures, further focussing on a subset that recurrently replicated across more than one replication method (see Fig. 3A) and/or more than one cancer type -- the ‘silver set’ of associations as defined above (Supplementary Table S7).

We noted that some associations in this set recurred in multiple methods/tissues, thus we introduced a more stringent tier of hits with a high priority for follow-up (‘golden set’). We focused on a subset that recurred in at least three tissues with effect size *d*>0.5 and significant at p<0.01 and at least once at FDR<25%. This resulted in 782 associations (alternatively a broader set of 4346 associations would be recovered, if not requiring strictly the same molecular target but instead permitting replication across different protein targets in the same pathway, while requiring 4 tissues; Supplementary Table S11 and S12). Validating this approach, we recovered a known association between the activity of the APOBEC mutagenic enzymes -- in this case, quantified via the SBS-APOBEC signatures SBS2 and SBS13 -- and sensitivity to ATR inhibitors^65,66^. The ATR inhibitors VE-822, AZD6738 and QL-VIII-58 were associated with SBS-APOBEC in lymphoma and in MSI colon cancer (at FDR<17%) with an additional suggestive result in head-and-neck cancer (p=0.004; Figure 5E). We suggest that, despite APOBEC activity in cell lines being intermittent^22^, the historical record of APOBEC activity captured in the mutational signatures is sufficiently predictive of vulnerabilities associated with APOBEC such as ATR inhibition, or – as we have recently shown experimentally and with a mutational signature analysis – loss of the abasic site sensor *HMCES*^67^.

A common occurrence among associations of the high-confidence set was involvement of signatures that were associated with DNA repair failures in previous analyses of tumor data. This included: MMR failures (the ‘summary signatures’ MMR1 and MMR2; see above, but also individual MMR SBS), BER failures (SBS36/18L/56L^5,68,69^, SBS30L/7bL/11L^4,70^), and replicative polymerase failures (particularly SBS14, SBS20 and SBS10b; additionally SBS56/10aL/36L may be in this group). As tentatively DNA repair associated signatures, here we additionally consider: SBS1 since one of its causes may be BER failures^12,71,72^, SBS12/26 based on the similarity of the spectrum to the “MMR2” signature^63^, SBS18/36L based on the similarity of the spectrum to the SBS36/18 and because it was found in MUTYH-variant patients^68,73^, and additionally SBS33L and SBS54 because they have strong indel components and are associated with sensitivity to *WRN* loss. Those DNA repair-associated signatures encompass 312 of 782 associations in the high-priority set, supporting the notion that DNA repair failures commonly result in drug vulnerabilities. We note that some role of DNA repair deficiencies is not excluded in some other signatures, e.g., SBS23/11L (which has 26 associations in the high-confidence set), because the constituent SBS23 and SBS11 both exhibit very strong transcriptional strand biases^12^ usually attributed to activity of transcription-coupled NER, and similarly so with SBS31 (which has 28 associations).

One prominent example (Supplementary Table S12) is that the signatures of DNA mismatch repair SBS15/6L and SBS20 as well as the summary signature SBS-MMR1 show multiple associations with sensitivity to histone deacetylase (HDAC) inhibitors (in colon, pancreas, skin lung, stomach/esophagus) as well as to inhibitors of HDAC-related sirtuin proteins (in colon); see Fig. 5B and Supplementary Figure 14A. The associations involve drugs such as belinostat, vorinostat, dacinostat and AR-42, which have broad specificity against many HDAC proteins and so are not highly informative about the specific targets necessary for sensitization. However, the gene knockout data from CRISPR screens recurrently implicated the *HDAC1* gene (Fig. 5B and Supplementary Figure 14A), while the drugs entinostat and CAY10603 as well as CRISPR editing converge onto *HDAC3* as likely targets. MMR-deficient colon cell lines and tumors are known to accrue mutations in the HDAC2 gene, which has a microsatellite repeat in its coding region^74,75^. This may provide a mechanism explaining associations in our data: a vulnerability may be created due to increased reliance on other *HDAC* genes, such as *HDAC1* and *3*. We suggest MMR mutational signatures may provide a useful genomic marker for applying HDACi. This has support in prior experiments on colorectal cell lines^76^ and in suggestive trends observed in a clinical study^77^ that reported a trend towards higher response rates to HDACi in MMR deficient colon cancer patients (odds ratios of ≈2 or higher for HDACi-containing regimens^77^). A caveat suggested by our data is that these effects may be HDAC gene specific and potentially tissue-specific. (Of note, the absence of associations that cross the statistical significance threshold in certain tissues does not necessarily imply that the association does not occur in those tissues, but could instead derive from differences in statistical power due to, for instance, tissues represented with more or with fewer cell lines; thus our analyses do not measure tissue-specificity). Additionally, some MMR signatures may be more predictive than others; we suggest SBS15, 6 or 20 as the strongest candidates for follow-up HDACi studies. Encouragingly, we did not find significant resistance associations involving HDACi and any MMR signature in colorectal, lung (adenocarcinoma or squamous cell carcinoma), pancreatic, gastroesophageal or skin cancer cells that replicated in genetic screening data. In addition to HDAC inhibitors, we highlight a possible vulnerability of MMR deficient cells exhibiting signatures SBS26/12L, SBS21 and SBS20, to nucleoside analogs (Supplementary Fig. 14B) Ara-G, fludarabine, nelarabine and cytarabine, as well as the antimetabolite methotrexate, in colon, brain, skin and pancreas at FDR<30% and further in bladder and myeloid leukemia at a suggestive threshold.

### Examples of other mutational signatures that predict drug sensitivity

In addition to MMR, we highlight an example involving a BER signature SBS36/18L/56L, associated with deficiencies in the MUTYH gene^5,68,71^ that processes oxidatively damaged nucleobases. This signature sensitized to cabozantinib in skin, and further (at permissive thresholds) in bladder and in pancreas (Fig. 5F). Cabozantinib is a tyrosine kinase inhibitor, typically considered to act by inhibiting VEGFR2, MET and RET, although it is recognized to inhibit many other kinases as well. Our replication analysis using CRISPR data suggests however that the selective activity of cabozantinib against SBS36-bearing, thus presumably BER-deficient cells, occurs via inhibition of AXL and/or KIT. Previous work suggests that inhibition of AXL suppresses DNA repair and sensitizes cells to various DNA damaging agents or repair inhibitors^78–80^ consistent with the hypothesis of a synthetic lethal association between BER loss and AXL inhibition.

Next, there were associations involving the common cancer drug doxorubicin and the signature SBS33L, which is dominated by the indel component (deletions) and has a minor contribution of single-nucleotide variants of the A>G type (Supplementary Figure 2); this signature also associates with WRN knockout sensitivity (Supplementary Figure 12) similarly as the known MMR-associated signatures. Sensitization to doxorubicin by SBS33L replicates in our ‘same-target’ test with mitoxantrone in colon and also in brain (at FDR 6% and 34%), and with epirubicin in pancreas (FDR=24%; Fig. 5D). This provides an example of a predominantly indel-based mutational signature that predicts response to intercalating agents in broad clinical use.

Finally, we suggest an example of a mutational signature that is according to our algorithm and data cell line specific, the SBS-CL5, that associates with sensitivity with multiple topoisomerase I inhibitors (including camptothecin, irinotecan/SN-38, and topotecan) and topoisomerase II inhibitors (etoposide, teniposide) across both GSDC and PRISM screening data in colon, with additional support for topoisomerase II inhibitor etoposide and *TOP2A* gene k.o. in skin (all at FDR<25%, Fig. 5C). This suggests that changes to DNA replication, repair or supporting activities acquired during cell culture might be used to elucidate mechanisms of sensitization to common anticancer drugs. Alternatively, it is possible that this SBS-CL5 profile rich in C>T changes (Supplementary Fig. 2) is a mixture of tumor SBS that were insufficiently resolved here (Supplementary Fig. 6) or conversely that it is a constituent component within some of the SBS found in tumors^12^ that were insufficiently resolved previously.

## Discussion

A classical way to treat tumors is to employ DNA damaging drugs and ionizing radiation to target the lessened or overwhelmed capacity for DNA repair in faster-dividing and/or replication stressed cancer cells, which may manifest as a mutator phenotype. We thus investigated the hypothesis that many mutational signatures can serve as markers of DNA repair failures targetable by drugs or by gene silencing. This generalizes over the known individual examples of mutational patterns stemming from deficient homologous recombination repair^15,17,18^ or mismatch repair, which suggested therapeutic strategies^58–60^. Cancer cell line panels that were systematically screened for drug responses^29,81^ and for gene loss effects^45,82^ provide a resource to test the hypothesis that mutational signatures are generally useful markers of drug response. However, the lack of matched healthy tissues from the same individuals means that methods to extract mutational signatures^22,23,33^ face a difficult challenge. The amount of residual germline variation, after filtering known variants from a population genomic database, still eclipses the amount of somatic variants. Even slight variations of germline mutational signatures between individuals and populations^25,27^ can strongly affect the trinucleotide context mutation tally of a cell line (or another tumor sample where a matched normal is not available). Our methodology addresses this issue, and further integrates indel features into signature inference; future refinements in the method as well as availability of whole genome sequences of cell lines will improve its accuracy, for instance permitting a more detailed set of indel descriptors, as applied in the recent tumor WGS mutation signature studies^12^.

Among those signatures we inferred from cell line exomes, the number of associations with drug response was comparable to that of other types of commonly used genomic markers that are based on driver mutations, copy number changes and promoter DNA methylation. Many such associations significantly replicated in independent data sets, including other drug screening methods and genetic screens, and thus mutational signatures appear to be similarly robust predictors as the commonly used, driver mutation-based or gene expression-based markers.

An important caveat is that the discovered associations do not necessarily imply a causal relationship: the mutational signature-generating process and the drug phenotype may be only indirectly associated. In other words, the mutational signature serves as a marker for another alteration, which is the proximal cause of the drug sensitivity or resistance. This is possible as with mutational signatures so with other genetic markers (mutations or copy number alterations in cancer genes) and also with gene expression markers. Because these various sets of features can be correlated across tumors, prioritizing the likely causal relations using statistical analyses is an important challenge ahead. Furthermore, another issue are false negatives – again similarly so with mutational signatures as with other markers – due to a small number of cell lines per tissue bearing a marker. Because drug sensitivity measurements can suffer from substantial noise resulting in less-than-ideal agreement between different screening data sets^34–36^, replication analyses across datasets like the one we performed can be conservatively biased.

Future work will refine the mutation signature markers and clarify the underlying mechanisms. One issue that merits attention is question is timing; a recent genomic analysis of cell lines^22^ has suggested that the activity of some mutational processes is intermittent. While a genome sequence reflects a record of mutagenic activity in the past, it may not always reflect current mutagenic activity, which is presumably more relevant for drug phenotypes. This factor is difficult to account for using bulk DNA sequencing from cell cultures (as the recent mutations would not rise to sufficient allele frequencies to be detected), and would bias our association analyses conservatively. This is because drug activity would presumably be more likely to reflect active mutational processes, which might not have had time to generate sufficient mutations. We note that a related issue could affect also the association analysis of drug activity with the more mainstream markers, individual mutations or CNA in cancer genes: a subsequent mutation may have happened that alters the effect of the original mutation via a genetic interaction. Given the rapid accumulation of genetic changes in cancer cell lines during culture^83,84^, it is well plausible that such events occur.

Another question regarding mutational signatures associated with drug activity across cell lines is that some of the signatures are thought to result not from DNA repair deficiencies but instead from exposure to mutagenic agents. Such signatures that recorded a high number of associations in our analyses (Fig. 4) for example SBS31, previously associated with pre-treatment with platinum drugs, or a SBS17a-like signature where SBS17 was associated with exposure to gastric juice since it was often observed in stomach and esophagus cancer, or SBS38 suggested to result from indirect DNA damage upon UV light exposure^12^. Because cell lines are presumably not exposed to these agents and thus the signatures are not ‘active’, it is intriguing that some drug sensitivity/resistance associations are found with such signatures (Fig. 4).

One possible explanation may be issues with signature inference methods, such that mutational signatures of different carcinogens are sufficiently similar that only the trinucleotide spectra do not easily unmix them. For instance SBS17 (mostly A>C) was proposed to result from varied mechanisms, some of which are from mutagenic chemotherapy exposure while others may be endogenous^85^, highlighting a possible example where current statistical approaches do not reliably deconvolve underlying biological mechanisms. Such issues with confounded mutational processes might be addressed by the use of additional features to derive mutational signatures, such as penta- or hepta-nucleotide contexts^87^, small indels^12^ and copy-number changes^13^.

Another explanation, speculatively, may be that a prior exposure to a carcinogen would select tumor cells with an altered DNA replication/repair state, which continues after the carcinogen is withdrawn, thus providing a vulnerability in cancer cells. For instance, therapy with the drug temozolomide (associated with SBS11) is known to select for MMR-deficient cells via loss of MSH6 function in brain tumors^88^. In other words, a temporary exposure to a carcinogen might prime the cells for successful treatment by certain drugs later. In our data, replicated drug associations involving the platinum drug-associated SBS31 were found (Figure 4). This suggests the exciting possibility that therapeutic exposure to a cancer drug might open up new treatment avenues after drug resistance arises, and that mutational signatures could provide markers to guide therapy of recurrent, drug-resistant tumors.

## Methods

### Human cancer cell lines and primary tumor data

We obtained WES bam files (human reference genome version hg19) of human cancer cell lines (n=1072; cancer cell lines from Genomics in Drug Sensitivity in Cancer (GDSC)) from European Genome-phenome Archive (EGA) (ID number: EGAD00001001039) and WES bam files (human reference genome version hg38) of primary tumors (n=6154) and their matched normal samples (n=6154) from the TCGA repository. We downloaded samples from the following TCGA cohorts: BLCA, BRCA, COAD, GBM, HNSC, KICH, KIRC, KIRP, LIHC, LUAD, LUSC, OV, PAAD, READ, STAD, THCA, UCEC. We aligned the human cancer cell line bam files to the human reference genome (version hg38) using the *bwa*^89^ software, and sorted and indexed them using the *samtools* software^90^. We used the *Strelka2* software (version 2.8.4)^91^ to call single nucleotide variants (SNVs) and small insertions and deletions. We called SNVs and indels for cell lines, primary tumors and normal samples. In samples where Strelka2 was unable to run, a re-alignment was performed using Picard tools (version 2.18.7)^92^ to convert the bams to FASTQ and, following that, the alignment was performed by executing bwa sampe (version 0.7.16a) with default parameters. The resulting bam files were sorted and indexed using Picard tools. We used SNVs and indels marked as “PASS” in the *Strelka2* output. We annotated SNVs and indels with minor allele frequencies (MAF) obtained from the gnomAD database^93^ (for SNVs and indels that could be found in GnomAD).

### Data for human cancer cell lines

We downloaded drug response data for 518drugs from GDSC (Release 8.3; June 2019)^29^. We used the natural logarithm of the 50% growth inhibition values (IC50) as a measure of the activity of a compound against a given cell line. If for the same drug activities were available from both GDSC1 and GDSC2 versions, we used GDSC1. We downloaded the information about drugs (the list of drugs’ putative targets and target pathways) from the GDSC website (https://www.cancerrxgene.org/). We manually curated the list to correct inconsistencies (Supplementary Table S13).

We obtained the following genetic features of cancer cell lines from the GDSC repository^21^: cancer driver genes (Muts), regions of recurrent focal copy number alterations (CNAs), hypermethylated informative CpG islands (HypMet), and microsatellite instability status (MSI). We downloaded ANOVA_input.txt files for 18 cancer types and pan-cancer analysis from https://www.cancerrxgene.org/gdsc1000/GDSC1000_WebResources/Pharmacogenomic_interactions.html. We downloaded cancer cell line gene expression data (“sanger1018_brainarray_ensemblgene_rma.txt”) from https://www.cancerrxgene.org/. We selected expression data for 1564 genes corresponding to the L1000 assay^30^ and to known drug targets^31^.

The drug response data (IC50) for 1,502 drugs was downloaded from PRISM^37^ database (secondary-screen-dose-response-curve-parameters.csv). Drugs and cell lines from PRISM and GDSC databases were matched via drug names, obtaining in total 348 cell lines and 178 drugs that overlap between the two databases.

The gene-level fitness effects for 16,827 genes in 786 cancer cell lines were downloaded as the Integrated cancer dependency dataset from Wellcome Sanger Institute (release 1) and Broad Institute (19Q3) (“integrated_Sanger_Broad_essentiality_matrices_20200402.zip “) from the Project SCORE database^45^ and matched to the GDSC cell lines via cell line names, obtaining in total 517 overlapping cell lines (https://score.depmap.sanger.ac.uk/downloads.

### Matching of cancer cell lines and primary tumors (ancestry matching procedure)

To ensure that genomic data is comparable between cell lines and tumors we performed the following steps: (i) we used only SNVs detected in the regions of the exome with well sequencing coverage (>=20 reads in 9 out of 10 randomly selected samples); (ii) to avoid gender bias during analysis, variants on X and Y chromosomes were not used; (iii) only uniquely mappable regions of the genome were used, as defined by the Umap k36 criterion^94^; (iv) we discarded regions with frequent somatic deletions, namely pan-cancer deletions of significant SCNA^95^ and frequently deleted chromosome arms (deleted in >18% of tumor samples)^96^; (v) we detected copy number changes of cell lines with CNVkit^97^ and removed deleted regions (log2 score < -0.3). From the remaining regions of the exome we selected common germline variants across the cell line exomes and the TCGA exomes (MAF>5% in GnomAD). To perform ‘ancestry matching’ of cancer cell lines with TCGA normal samples, we performed Principal Component Analysis (PCA) on the matrix of common germline variants, followed by clustering according to principal components (PC) (see below).

#### Clustering of cell lines and TCGA germline samples

We employed robust clustering (*tclust* algorithm^98^; discards outlying samples) on the first 140 principal components derived from the common germline variants. Initially, we considered the first 150 principal components, however, some PCs were attributed to the batch effect, i.e., they separate cell lines from TCGA samples. We removed the top 10 such PCs as determined by feature importance of the random forest classifier (*randomForestSRC* R package) trained to distinguish between TCGA samples and cell lines. We used the remaining 140 PCs as an input to the tclust algorithm. Outlying samples, as determined by tclust, were discarded. We varied the number of clusters from 4 to 20, determining the optimal number of clusters (13) by using simulated cell line exomes (Figure 2B, C; see below). The cancer cell lines were matched to their ‘ancestry-matched’ TCGA normal samples, i.e., those belonging to the same cluster as the cell line. We then used ‘ancestors’ to augment the filtering of germline SNVs from cancer cell line exomes, in addition to filtering according to the common practices to retain only variants absent or present in very low frequencies in population databases (see below).

#### Filtering of germline SNVs from cell lines

Here we considered SNVs from the regions of the exome as specified before (steps i-v), with the difference that we used SNVs with the sequencing coverage of >=8 reads in at least 90% samples (9 out of 10 randomly selected samples). We next filtered germline variants from cancer cell lines following common practices^20,22,81^ -- here, removing population variants (found at MAF>0.001% in the gnomAD population database); additionally we filtered germline variants that appeared >5% of samples in our TCGA data set or cell lines data set which removes germline variants that might be particular to this data set and also suspected sequencing artefacts). Next, from the remaining SNVs, for each cell line and TCGA germline sample, we calculated its trinucleotide mutation spectrum. The mutation spectrum contains 96 components, which are the counts of six possible mutation types (C>T, C>A, C>G, A>T, A>G and A>C, considered DNA strand-symmetrically) in 16 possible 5’ and 3’ neighboring nucleotide contexts for each mutation type. For each cell line, from the cell line’s 96 trinucleotide spectrum, we subtracted the median 96 trinucleotide frequencies of its TCGA ‘ancestry-matched samples’ (i.e., the TCGA normal samples belonging to the same population cluster as the cell line in the PC analysis of common germline variants, see above). In case the subtraction resulted in a negative count for some of the context, we set it to zero.

#### Insertion and deletion types

In addition to the standard 96 trinucleotide spectra, to extract mutational signatures from cell lines (see below) we considered additional features based on small insertions and deletions. We filtered the regions of the exome using the steps (i-iv) as described above. Similarly as for SNVs, we discarded indels found at MAF>0.001% in the gnomAD population database and filtered the ones that appeared in >5% of cell line samples. The remaining indels were classified into 4, 5, 8, 14 or 32 different indel types types considering: the length of insertion or deletion (considering lengths 1, 2, 3-4, 5+), microhomology at deletion sites (considering the lengths of microhomology 1 or 2+), and microsatellites at indel loci (considering repeat sizes of 1 and 2-5+). We benchmarked the cell line mutational signatures (see below) that used different indel types. The most favorable indel types according to that benchmark were the simplest where we differentiate 4 indel types: deletions with microhomology (Del-MH), deletions at microsatellite loci (Del-MS), other deletions (Del-Other), and insertions at any locus (Insertion).

### Simulated cell line exomes

We used TCGA exomes to simulate cell line exomes (or more precisely, the variant calls that would originate from a cell line exome) in order to benchmark our ancestry matching procedure for efficacy in reconstructing the mutation spectrum of cancer cell line exomes. Additionally we used the benchmark to optimize the number of clusters in the inference of subpopulations. To simulate variant calls from cell line exomes, we performed the variant calling for tumor samples in the same way as for the cell lines, i.e., without matched normal samples. In addition, for every sample we merged SNV calls obtained thus with the somatic SNV calls of tumors, ensuring that the true somatic mutations are a part of a simulated cell line to enable an accurate estimation of errors. In the tumor types that were represented both in the set of cancer cell lines and the set of TCGA tumors, we randomly selected 450 tumor samples taking approximately the same number of samples per tumor type as in cell lines. In the subsequent analysis involving simulated cell line exomes, we removed their corresponding normal samples from the pool of germline samples. We performed the ancestry matching procedure involving simulated cell line exomes in the same way as described above (Figure 1E) and evaluated the accuracy of mutation spectrum reconstruction by comparing the mutation spectrum of a tumor (ground truth) to the corresponding simulated cell line mutation spectrum obtained by ancestry matching. As an accuracy measure we used Root Mean Squared Error (RMSE). To estimate the optimal number of subpopulations we varied the number of clusters in the tclust algorithm from 2 to 20, with 13 being the optimal according to the average RMSE of 450 simulated cell lines (Fig. 1C).

To investigate the variation between germline mutational spectra across human (sub)populations (as reported recently^25^), we performed hierarchical clustering (based on cosine similarity; *stats* package in R) of the median mutation spectra of TCGA subpopulations (i.e., median mutation spectra of the TCGA normal samples within clusters) (Fig. 2C). In addition, we performed PC analysis of mutation spectra of TCGA normal samples showing that the main principal component separates the main ethnicity groups (Fig. 2D,E). We note that, in the previous steps, clustering to infer subpopulations was performed *via* common variants (see above), and the trinucleotide spectra were determined after excluding the non-rare variants (MAF>0.001% in the gnomAD population database and variants that appear in >5% of samples in our TCGA data set) ruling out circularity.

Next, we compared our ancestry matching procedure (involving 13 clusters) to the baseline method of determining somatic variants in a cell line genome (i.e., filtering of germline variants according to MAF where SNVs with MAF>0.001% and that appeared in >5% of samples in our dataset are removed) and the bootstrap self-similarity method (Fig. 1D). The bootstrap self-similarity error was calculated as an error between true and randomly perturbed somatic mutation spectrum, averaged over 100 random runs. For random perturbation we used the *sampling* function of the *UPmultinomial* R package.

### Extraction of mutational signatures

We extracted cancer cell line mutational signatures from 96 component trinucleotide mutation spectra of cancer cell lines obtained with the ancestry matching procedure. In total we used mutation spectra of 930 cancer cell line exomes (Supplementary Table S14). Note that some cell lines were excluded because they were assigned to the outlier cluster of the *tclust* algorithm, based on their common germline variants (see above). To extract mutational signatures we used a custom R script based on non-negative matrix factorization (NMF) as described by Alexandrov *et al*.^24^. We additionally implemented a number of signature extraction procedures proposed in the literature recently^12,33,63^ and benchmarked the resulting signatures according to the several criteria: (a) The agreement with an established set of SBS signatures (PCAWG signatures^12^) measured as the number of PCAWG signatures that were recapitulated at >=0.85 cosine similarity cutoff; (b) The similarity of the cell line signature exposure profiles across cancer types to the exposure profiles of PCAWG signatures, measured as a cosine similarity between an average exposure per-cancer-type profile of a cell line signature and its matching PCAWG signature (for signatures recapitulated at >=0.85 cosine similarity); (c) The accuracy of the signatures in predicting drug sensitivity profiles across cancer cell line panels (see below for method).

From the matrix containing mutation spectra of samples, we generated 300 matrices by bootstrap resampling. One bootstrap sample is obtained by applying the sampling function of the *UPmultinomial* R package to each sample’s spectrum. Next, we applied the NMF algorithm to each of the bootstrap samples (nmf function of the *nmfgpu4R* R package to obtain different NMF runs; we used the Multiplicative update rules algorithm^99^ with 10000 as the maximal number of iterations). For each bootstrap sample, we varied the number of signatures from 2 to 40. Computations were performed on an NVIDIA GeForce RTX 2080 Ti graphics card.

We implemented the following modifications to the above basic procedure (each tested independently and some in combinations; Supplementary Table S1) to obtain the candidate cell line signatures which were matched to the known tumor PCAWG signatures (see below): (i) Similarly as done in the seminal work of Alexandrov *et al*.^24^, we clustered each batch of NMF solutions and used the cluster medoids as candidate cell line signatures. We used the *clara* function in clusters R package, with Euclidean distance and standardization and “pamLike” options; the number of samples to be drawn from the dataset was set to 10% of the number of samples. Each batch of NMF solutions obtained as described before (one batch corresponds to 300 x n solutions, where n is the number of signatures; varies from 2 to 40) was clustered into k clusters with k-means clustering, varying k from 2 to 30. (ii) We applied the RTOL-filtering (relative tolerance) as proposed by Degasperi *et al*.^63^, where (separately for each different number of signatures parameter) NMF runs that diverge for more than 0.1% from the best NMF run are removed, as measured by the root-mean-square deviation of the factorization. (iii) We implemented the ‘hierarchical extraction’ procedure proposed by Alexandrov *et al*.^12^ where NMF is repeated iteratively while removing (or down weighting) the well-reconstructed samples (cosine similarity above a specified threshold) from a previous iteration to uncover new signatures. We allowed a maximum of 3 iterations and tested different cosine similarity thresholds ranging from 0.95 to 0.99. In the case of down weighting, we multiplied sample’s mutation spectra by 0.05 instead of removing it. (iv) Similarly as in Ghandi *et al*.^33^, we performed joint signatures inference of cell line exomes and tumor exomes. To the initial matrix containing mutation spectra of cell lines, we added mutation spectra of 7755 TCGA somatic exomes which were preprocessed in the same way as cell line exomes: the same regions of the exome were used (see above for filtering by coverage, mappability, etc.).

From the candidate cancer cell line mutational signatures obtained by the above described methodologies, we searched for the signatures that closely resemble the ones that were previously found in human cancers^12^ (referred to as PCAWG signatures in the text). We compared the individual signatures to the PCAWG signatures searching for the closest matching cell line signature for each PCAWG signature. As a final set of tumor-like cell line signatures we kept the closest matching cell line signature if its cosine similarity to the best matching PCAWG signature exceeded 0.85. For cell line signatures that used indel features in addition to 96 tri-nucleotide spectra, cosine similarity was calculated only on 96 spectra. We considered an additional criterion for matching of cell line and tumor signatures where in addition to the mutation-spectrum cosine similarity, a cosine similarity between signature’s average per-cancer-type exposure profile (for cancer types available in both cell line and PCAWG data). The spectrum-cosine and exposure-cosine similarities were combined into a single metric given the weight ‘w’ controlling the relative contribution of each: w×spectrum-cosine + (1-w)×tissue-cosine. We tested different weights: w=0.5, 0.7, 0.9, and 0.99. For each PCAWG signature, from a set of cell lines signatures that match it with spectrum-cosine>=0.85, we selected the best matching cell line signature as the one with the highest combined metric.

We considered using an additional regression post-processing for assigning signature exposures to samples, similarly as done by Petljak *et al*.^22^ and Alexandrov *et al*.^12^ We used the sigproSS tool of SigProfilerExtractor framework^100^ to attribute exposures to the extracted cell line signatures (i.e., signatures were used as an input to sigproSS). Additionally, we used a custom script based on regularized regression from glmnet R package enforcing different degrees of sparseness. We considered Ridge, Lasso and Elastic Net regression to model cell line’s mutation spectra as a combination of cell line signatures, where the resulting regression coefficients are considered as exposures to signatures. We enforced non-negative coefficients, no interaction term, and used crossvalidation to determine the optimal value of the lambda parameter.

The different procedures for mutational signature extraction we implemented were evaluated according to the above described criteria (a-c), results are presented in Supplementary Table S1. As a final set of signatures we chose signatures obtained with hierarchical extraction (using 0.97 cosine similarity threshold for sample removal) and using 4 indel features in addition to 96 trinucleotide types. No post-processing of exposures to signatures was performed since it did not yield improvements according to the evaluation criteria, thus, we used the raw NMF scores of the exposure matrix.

This procedure yielded 50 signatures (Supplementary Fig. 2). The cell line signatures are named according to the PCAWG signatures they resemble, e.g., the cell line signature name SBS46/26/12L denotes that for PCAWG SBS46 this signature was the closest match (i.e., SBS46 is the primary signature); 26/12 denotes that the signature also resembled PCAWG signatures SBS26 and SBS12 (at cosine similarity > 0.85). The suffix “L” (for “like”) denotes 0.85 ≤ cosine similarity < 0.95 (a somewhat less-close match), while the absence of the suffix “L” means cosine similarity ≥ 0.95. Names of signatures other than the primary signature (if present) are ordered by decreasing cosine similarity. A notable modification from the signature extraction method presented in Alexandrov *et al*.^24^ is that we did not limit to a single value of the ‘number of signatures’ NMF parameter (choosing based on measures of fit and of consistency, as in Alexandrov *et al*.^24^). Instead inference was run for many values of this parameter, and the final set of mutational signatures consists of solutions from different values for the ‘number of signatures’ parameter.

We next searched for cell line-specific signatures, i.e., signatures that commonly appear in cell line data and do not resemble any of the known tumor signatures. To this end, we employed k-means clustering (*clara* function in *clusters* R package, with Euclidean distance, standardization, “pamLike” options and 10% as the number of samples to be drawn from the dataset). Each batch of NMF solutions (from the signature extraction method selected as final by the evaluation) was clustered into *k* clusters with k-means clustering varying k from 2 to 40. We chose the clustering result (i.e., a set of signatures) where the agreement with PCAWG signatures was maximized in terms of the number of cluster medoids that resemble PCAWG signatures (at cosine similarity >0.85). From such a set of signatures we selected the ones dissimilar from any of PCAWG signatures (cosine similarity <0.8), yielding in total 5 cancer cell line specific signatures (named SBS-CL). These SBS-CL signatures appear together with real signatures and are robust (i.e., they are cluster medoids) therefore they are also likely bona fide mutational signatures which might originate, for instance, from cell-line specific mutational processes.

In total, this yielded 55 cancer cell line signatures (50 corresponding to known tumor signatures, and 5 additional cell line specific signatures) (Supplementary Fig. 2). To investigate if some of the extracted signatures are a result of incomplete separation of other signatures we considered the coefficients of the Lasso regression (*glmnet* R package) (Supplementary Fig. 6), where we modelled cell line signatures extracted in this work as a linear combination of PCAWG signatures (Supplementary Fig. 6).

### Predicting drug response

We built predictive models of drug response using the Random Forest (RF) algorithm as implemented in the ‘randomForestSRC’ R package. We compared the predictive performance of different predictors: mutations in cancer driver genes (Muts), recurrent copy number alterations (CNAs), DNA hypermethylation (HypMet), gene expression, previously reported cancer cell line mutational signatures^22,23,33^, and mutational signatures extracted in this work. The dependent variable was a response to a drug; we iterated this over all drugs. We built cancer type specific models for each drug separately, considering only models where at least 15 cell lines with drug response were available to train the model. For model validation we used 10-fold cross validation which was repeated five times to get more stable results. We built random forests with 100 trees and a minimal number of samples in terminal nodes set to 2. We assessed the predictive performance of the predictors by the relative root-mean-square-error (RRMSE), i.e., a root-mean-square-error divided by the root-mean-square error of the default model. The default model predicts a constant value for all cell lines equal to the average ln IC50 across the training set. RRMSE < 1, therefore, denotes better accuracy better than the one of the (uninformative) default model. We define such models as predictive. Results are presented as an average RRMSE per cancer type across all models built for a cancer type (Figure 3A). Note that, all drugs were not necessarily tested exhaustively across all cancer cell lines, therefore, the number of models per cancer type may differ (one model corresponds to one drug) due to missing data (<15 cell lines with drug response data available), as well as the number of model per different predictors (mutational signatures, gene expression, oncogenic mutations, copy number alterations, DNA hypermethylation) due to data availability. In the analysis of complementarity between different predictor types (Fig. 2D, E, F; Supplementary Fig. 7B) we thus report relative numbers of predictive models to facilitate fair comparison.

To assess the statistical significance of the differences in the predictive performance (Figures 3B and 3C), we follow the recommendations given by Demšar^101^. More specifically, to statistically compare the predictive performance of multiple predictors over multiple datasets we use the corrected Friedman test and the post-hoc Nemenyi test^102^. Here, a dataset corresponds to a pair of one cancer-type and one drug. Due to the requirements of this test, only the intersection of drugs modelled across all cancer types and predictors were considered (due to missing data the number of drugs modelled per cancer type and/or predictor may differ; see above). For each drug-cancer-type pair, predictors are ranked according to their RRMSE (for that drug-cancer-type pair), where rank 1 corresponds to the best (i.e.,the lowest) RRMSE, 2 to 2nd best, etc. Ranks are then averaged across all drug-cancer-type pairs to obtain the average rankings of predictors. The Nemenyi test performs pairwise comparison of predictors’ performance based on the absolute difference of the average rankings of the predictors to assess whether there are statistically significant differences among predictors. The test determines the critical difference (CD) for a given significance level α, if the difference between the average ranks of two predictors is greater than CD, the null hypothesis (that the predictors have the same performance) is rejected., i.e., there is a statistically significant difference between the two predictors. The results from the Nemenyi post-hoc test are presented with an average ranks diagram^101^. The average predictor rankings are depicted on an axis, in such a manner that the best ranking algorithms are at the left-most side of the diagram. The algorithms that do not differ significantly (in performance) for a significance level of 0.05 are connected with a bold line, therefore, predictors that are not connected are statistically significantly different according to the test

### Associations with drug response

#### Associations with continuous values of the mutational signature exposures

We searched for associations between ‘exposures’ to cancer cell line mutational signatures (signature weights from the NMF inference; Supplementary Table S3) and response to drugs (ln IC50). For each cancer cell line, we normalized the exposures to get relative signature contributions: exposure to each signature was divided by the sum of exposures across all signatures for a cell line. In addition to mutational signatures, we considered associations with genetic and epigenetic features of cell lines as defined previously^21^: mutations in cancer driver genes (Muts), regions of recurrent focal copy number alterations (CNAs), and hypermethylated CpG islands (HypMet).

Association analysis was performed similarly as in Iorio *et al*.^21^. For each drug, a linear model was constructed (*stats* package in R) where the dependent variable (i.e., drug response) was represented as a combination of cancer type, the growth medium type used for the cell line, and values of a feature (i.e., exposures to a mutational signature, or a status of a genetic feature). For each available feature-drug pair, we constructed cancer-type specific predictive models. Prior to the association search, we removed 21 cell lines that exhibited either sensitivity or resistance nonspecifically towards a large number of drugs (15 cell lines reported by Abbas-Aghababazadeh *et al*.^103^ and 6 outlier cell lines considering median ln IC50), as well as additional 29 cell lines that were reported to be misclassified^104^. In addition, for each cancer type we removed outlier cell lines by the total number of mutations (remaining after the filtering with the ‘ancestry matching’ procedure; see above), here defined as having the number of mutations larger than 3 X interquartile range + upper quartile, or lower than 3 X interquartile range - lower quartile (calculated for each cancer type separately). We considered only models where mutational signature or genetic feature had at least three non-zero values.

P-values of linear regression models were obtained through type-II ANOVA modelling and were adjusted for multiple testing with the Tibshirani-Storey method^38^. A condition for statistical significance was set to false discovery rate (FDR)<25% and p-value<0.001.

#### Association that replicate in independent datasets

We performed two-way association testing where we searched for robust associations that replicate in two independent datasets. We considered three different types of ‘two-way’ tests where we enforce that, for a given drug, an association between a particular feature (a mutational signature or a genetic feature) and the drug response from the GDSC database is replicated: (1) in the PRISM drug screening database (GDSC/PRISM test) for the same drug, or (2) with another drug from the GDSC database that share the same molecular target (GDSC/GDSC (same target) test), or (3) in the Project SCORE CRISPR/Cas9 genetic screen as an association with a protein-coding gene fitness score of one of the drug’s target proteins (GDSC/PSCORE test).

We considered cancer-type specific associations. We required that in both tests of a ‘two-way’ test an association is detected in the same cancer type. We merged some similar cancer types with a small number of cell lines: esophagus carcinoma and stomach adenocarcinoma (denoted as ESCA_STAD), glioblastoma and brain lower grade glioma (denoted as GBM_LGG), and head and neck squamous cell carcinoma and lung squamous cell carcinoma (denoted as HNSC_LUSC). We additionally considered two groups of cell lines obtained by dividing colorectal cell lines according to microsatellite instability status (denoted as COREAD_MSI and COREAD_MSS; other cancer types did not have enough cell lines with microsatellite (in)stability labels to warrant such division). Outlier cell lines were removed in the same way as described above. We used binarized exposures to mutational signatures where, for each signature, values below the 10th percentile were set to 0, and the rest to 1 (41 out of 44 signatures had more than 10% of zero values, therefore, the resulting binarized signatures also often have more than 10% zeros in practice) (Supplementary Table S4). We used normalized (relative) signature exposures, as described above. We considered only tests with at least 8 cancer cell lines and at least 3 non-zero values of a feature.

All of the association tests were performed by modelling the drug response (or gene fitness score) to associate it with the status of a feature (i.e., a mutational signature or a genetic feature) searching separately for sensitivity and resistance associations. For each association, we calculated the association score as the minimum (in the case of sensitivity) or maximum (resistance) effect size (Cohen’s d) of the two independent datasets (i.e., positive Cohen’s d implies sensitivity and negative resistance associations). Here, effect size is the Cohen’s d statistic: difference of mean drug sensitivity (ln IC50) between the cell lines having the feature and those not having it, divided by the pooled standard deviation of the data. To obtain the association’s empirical p-value, we performed a randomization procedure where we calculated the association score 50000 times for the randomly shuffled features. For the sensitivity test, the formula for the pvalue is: 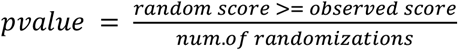and for the resistance is: 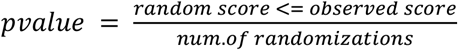. Due to the computational burden, we performed the randomization procedure only for associations that had an effect size >0.2 in the primary test. The empirical p-values were adjusted with the Tibshirani-Storey method^38^.

We considered aggregated mis-match and APOBEC mutational signatures where we merged p-values and effect sizes of SBS6, SBS15/6L, and SBS44L (denotes as SBS-MMR1); SBS21 and SBS26/12L (denoted as SBS-MMR2); and SBS2 and SBS13 (denoted as SBS-APOBEC). Pooled p-values of aggregated signatures were obtained by Fisher’s method, while pooled effect size was obtained by averaging. Pooled p-values were adjusted the same as described above.

We consider a two-way association as statistically significant if FDR<25% and additionally we imposed an effect size threshold Cohen’s d>0.5 in both tests (based on a known set of positive control associations (Supplementary Fig. 10). For some analyses we considered an additional set of associations with an unadjusted p-value threshold of < 0.01 and the same effect size threshold of Cohen’s d > 0.5. As a rule-of-thumb for interpretation, Cohen’s d = 0.2, 0.5 and 0.8 correspond to small, medium and large effect sizes, respectively^105^. We used QCEWAS R package to calculate the lambda score to estimate the inflation of p-values (calculated separately for sensitivity and resistance associations).

#### ‘Golden’ and ‘Silver’ sets of high-priority associations

We collated a list of 13846 associations (the ‘silver set’; Supplementary Table S7). To make this list, we considered all associations tested in the three ‘two-way’ replication tests that pass the permissive criterion of significance (effect size *d*>0.5 in both tests and a nominal *p*<0.01). From these, an association (between a feature and drug within a certain cancer type) was listed in the silver set if it was confirmed in more than one ‘two-way’ replication tests, or was seen in more than one cancer type, or if an association was seen with another drug that targets the same pathway. We require that at least one of the supporting associations has FDR<25%. Additionally, we collated a ‘golden set’ of high-priority list of associations involving mutational signatures which we consider to be suitable for follow up work: we require that and association involving the same drug is supported in at least three cancer types where at least one association has FDR<25% (782 associations; Supplementary Table S11) or that an association involving drugs targeting the same pathway is supported in at least 4 cancer types where at least two associations have FDR<25% (4346 association; Supplementary Table S12).

## Abbreviations of cancer types

A list of abbreviations of cancer types used in this study:

ALL/CLL: Acute/chronic lymphoblastic leukemia;
BLCA: Bladder urothelial carcinoma;
BONE: Bone cancer other/not classified further;
COREAD: Colon adenocarcinoma and rectum adenocarcinoma;
COREAD_MSI: microsatellite instable colon and rectum adenocarcinoma;
COREAD_MSS: microsatellite stable colon and rectum adenocarcinoma;
ESCA: Esophageal carcinoma;
ESCA_STAD: esophagus carcinoma and stomach adenocarcinoma;
EWING: Ewing’s sarcoma ;
GBM: Glioblastoma multiforme;
GBM_LGG: glioblastoma and brain lower grade glioma;
HNSC: Head and neck squamous cell carcinoma;
HNSC_LUSC: head and neck squamous cell carcinoma and lung squamous cell carcinoma;
KIRC: Kidney renal clear cell carcinoma;
LAML: Acute myeloid leukemia;
LGG: Brain Lower Grade Glioma;
LIHC: Liver hepatocellular carcinoma;
LUAD: Lung adenocarcinoma;
LUSC: Lung squamous cell carcinoma;
LYMP: Lymphoma;
MESO: Mesothelioma;
MM: Multiple myeloma;
NB: Neuroblastoma;
OV: Ovarian serous cystadenocarcinoma;
PAAD: Pancreatic adenocarcinoma;
SARC: Sarcoma other/not classified further;
SCLC: Small cell lung cancer;
SKCM: Skin cutaneous melanoma;
STAD: Stomach adenocarcinoma;
THCA: Thyroid carcinoma;
UCEC: Uterine corpus endometrial carcinoma.

## Supplementary figures

**Supplementary Figure 1.**
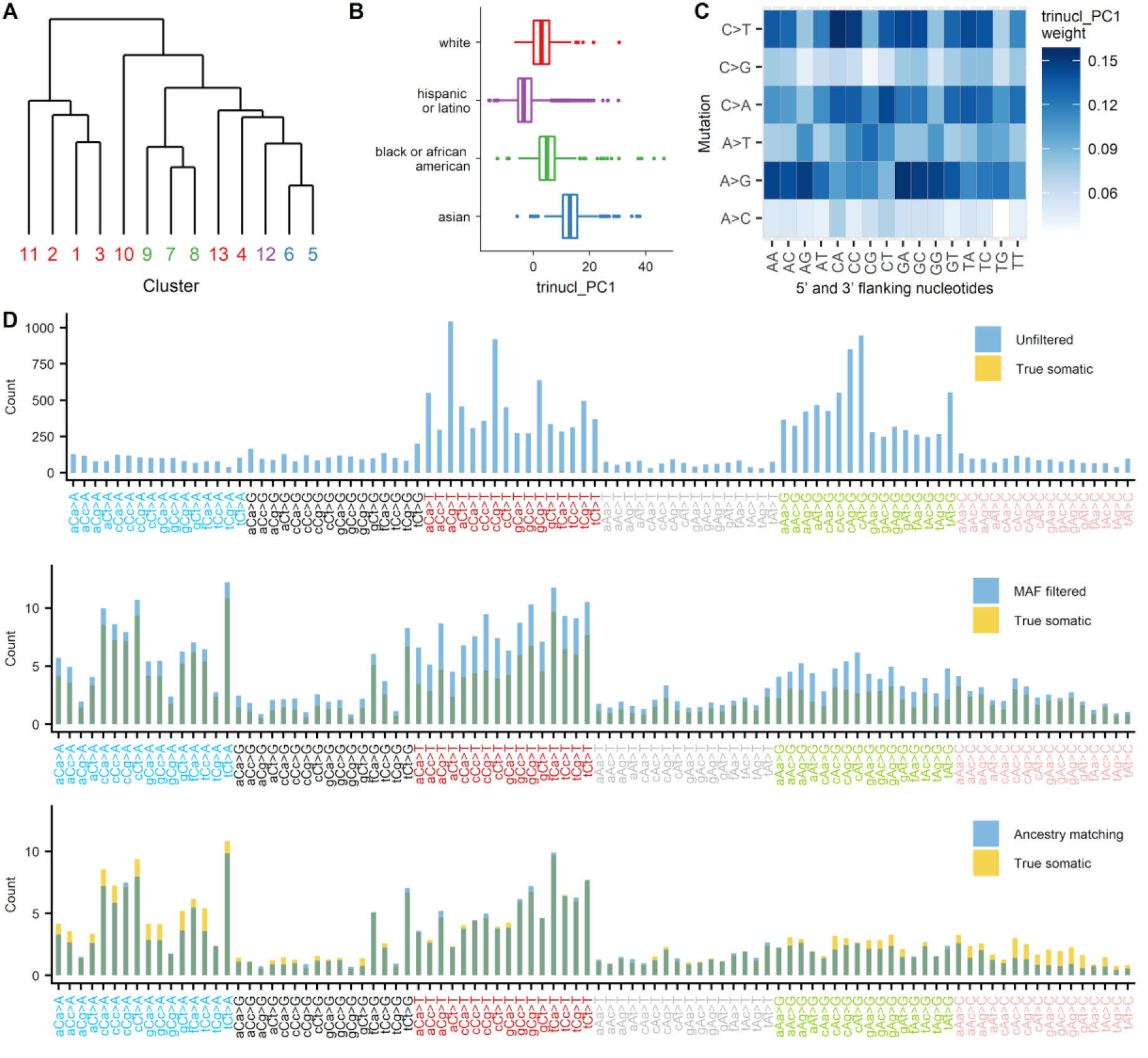
Ancestry signal in cell line exomes is correlated with trinucleotide mutation spectra of rare germline variants, which can confound mutation signature analysis. (A) Hierarchical clustering of the median trinucleotide spectra (derived from rare SNVs, MAF<0.001%, thus independent of the PC analysis) of the 13 ethnicity subgroups procedure shows that the intra-ethnicity tri-nucleotide profiles bear more similarity to each other than intra-ethnicity profiles (the subgroup labels are colored by the prevalent ethnicity within the cluster, with colors corresponding to panel B). (B, C) The main component of the PC analysis of tri-nucleotide spectra of the very rare germline variants (i.e., those that often cannot be removed by filtering using population databases; here MAF<0.001% or absent in gnomAD) separates the major ethnicity groups. (D) Average 96 trinucleotide spectra of 450 simulated cell lines. True somatic mutations are overlaid with the MAF unfiltered spectra, spectra obtained by the commonly used approach of filtering (MAF<0.001%) according to the population databases (upper panel) and the spectra obtained with the ‘ancestry matching’ method (lower panel).

**Supplementary Figure 2.**
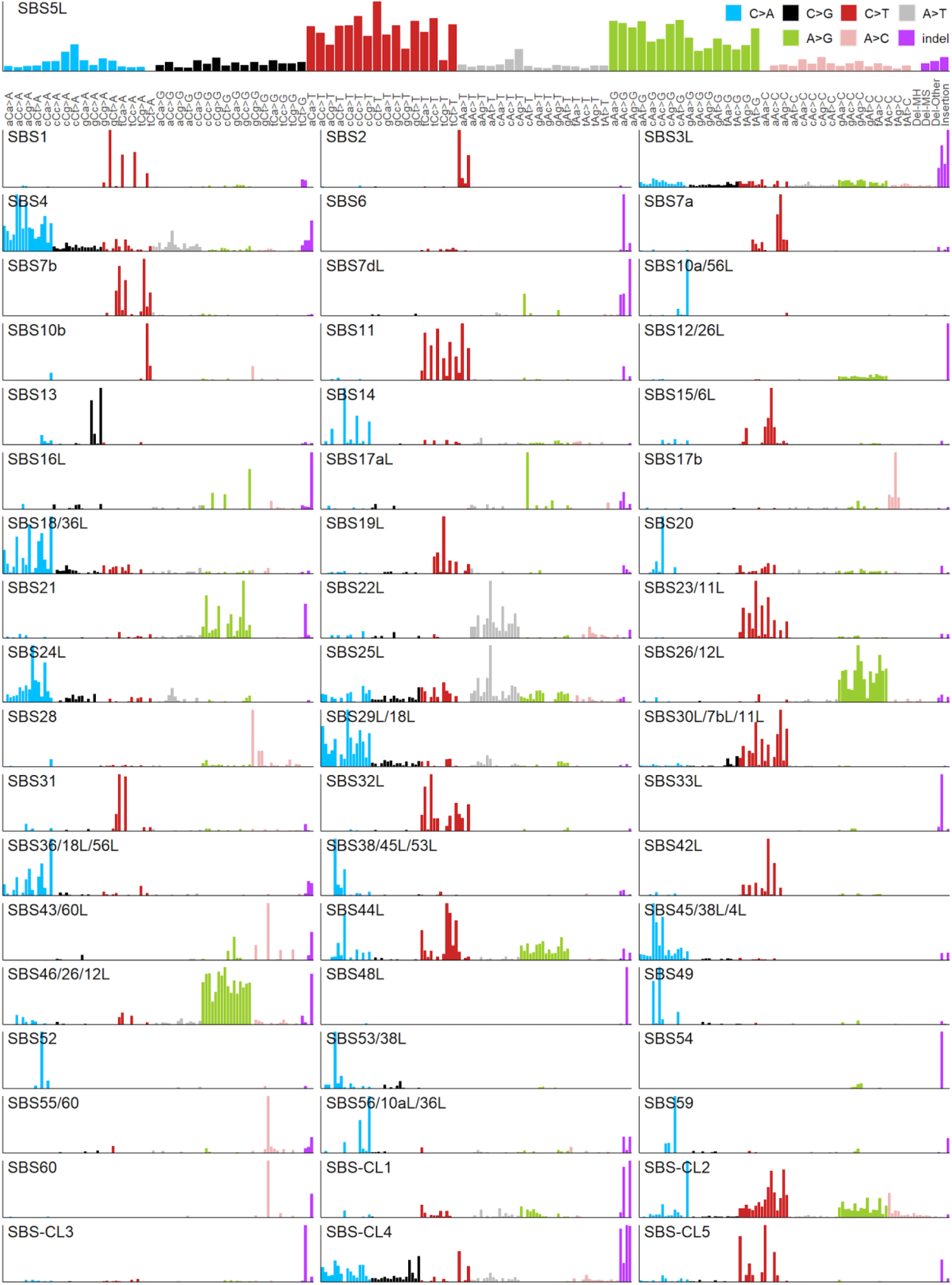
The 96 tri-nucleotide and 4 indel-type profiles of the cell line signatures inferred in this study. “Del-MH” are deletions with microhomology, “Del-MS” are deletions at microsatellite loci, “Del-Other” are other deletions, while “Insertion” are insertions at any locus. The y-axis corresponds to NMF weights.

**Supplementary Figure 3.**
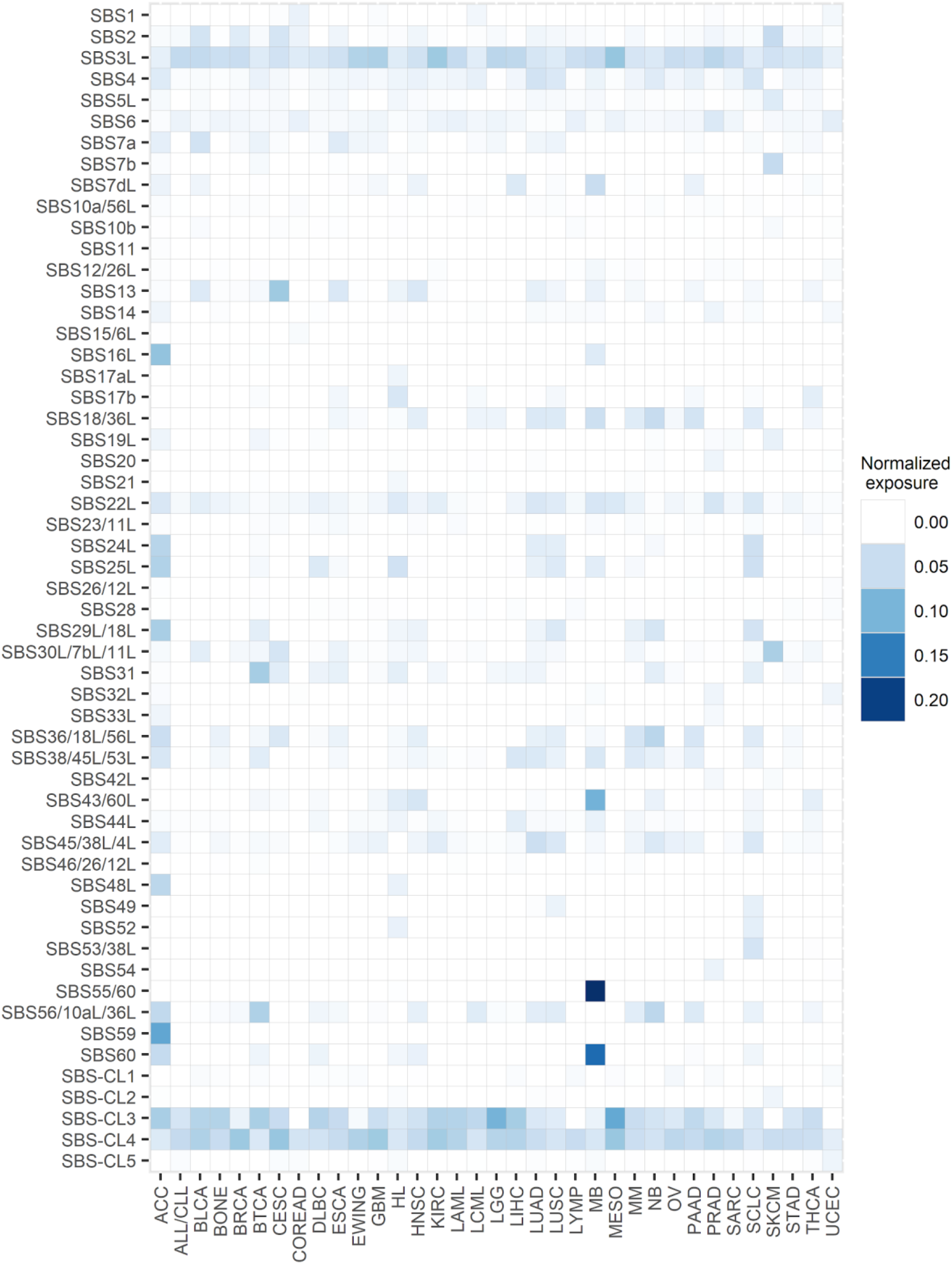
Estimated exposures of tissues-of-origin of the cancer cell lines to mutational signatures inferred in this work. The color intensity in the heatmap corresponds to the median NMF score per cancer type. For each cell line, the scores were normalized to represent the relative contribution of each signature in the exome of that cell line.

**Supplementary Figure 4.**
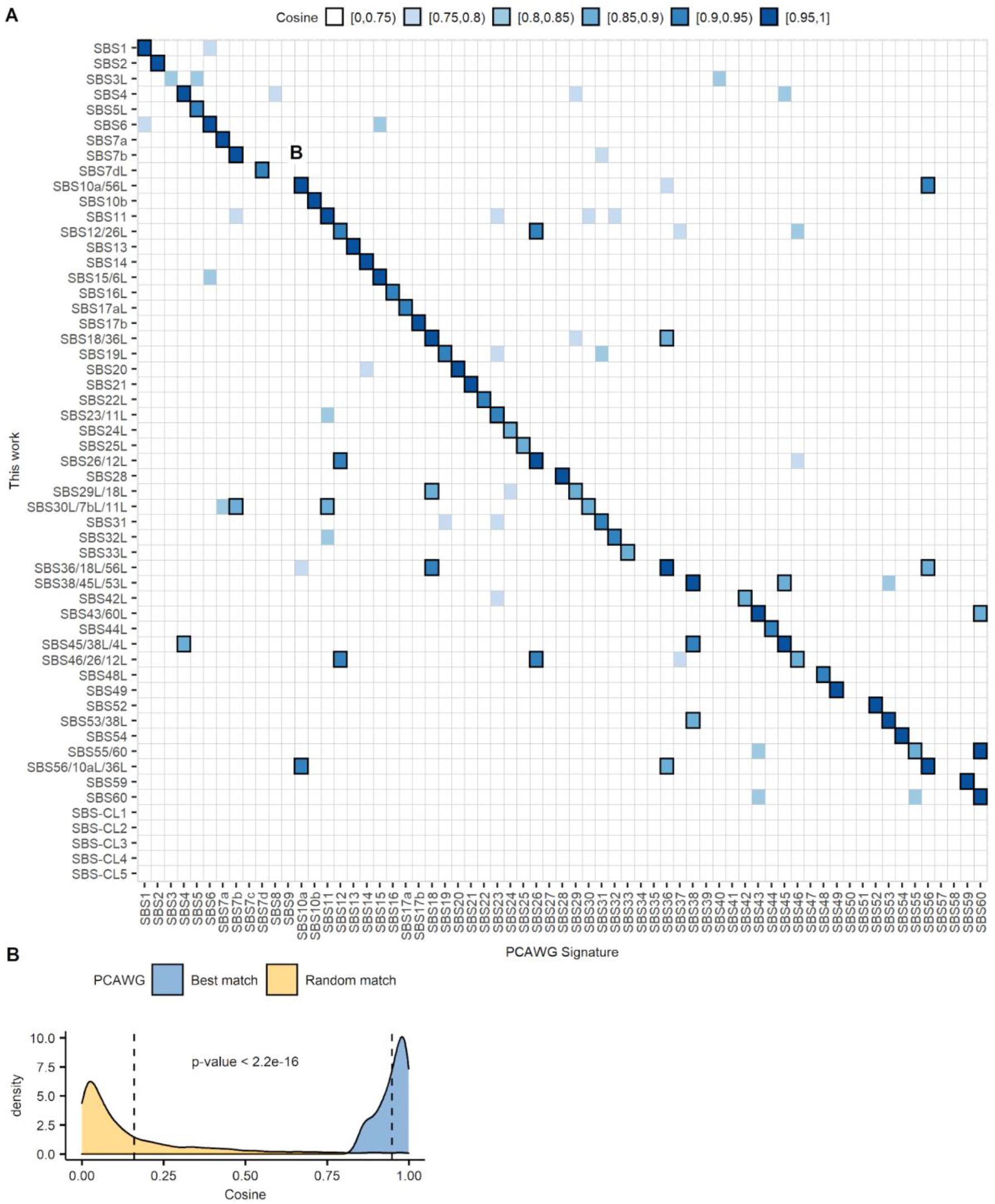
Cell line mutational signatures sometimes map to multiple tumor mutational signatures. (A) Cosine similarity between cell line mutational signatures and known tumor mutational signatures^12^. Some of the cell line signatures discovered in this work are highly similar (cosine>0.85) to more than one tumor signature (marked with bold black border rectangles). Such ambiguity is made clear in the naming convention of the cell line signatures where all of the highly resembling tumor signatures are included in the name of the cell line signature. Signatures that closely match previous tumor signatures (cosine similarity >= 0.95) are denoted with a corresponding name of the SBS tumor signature^12^, the suffix “L” (for like) denotes a less close match (0.85 <= cosine similarity < 0.95). Cancer cell line specific signatures (denoted as SBS-CL) are dissimilar to any of the tumor signatures and may represent cell-line specific mutational processes, germline mutations, or artefacts. (B) The density plot of cosine similarities of best matches of each of our cell line signatures to the PCAWG signatures, contrasted against a randomized baseline where each of our signatures is randomly matched to a PCAWG signature (repeated 100 times). The p-value of Mann-Whitney test comparing the means of the two distributions is shown.

**Supplementary Figure 5.**
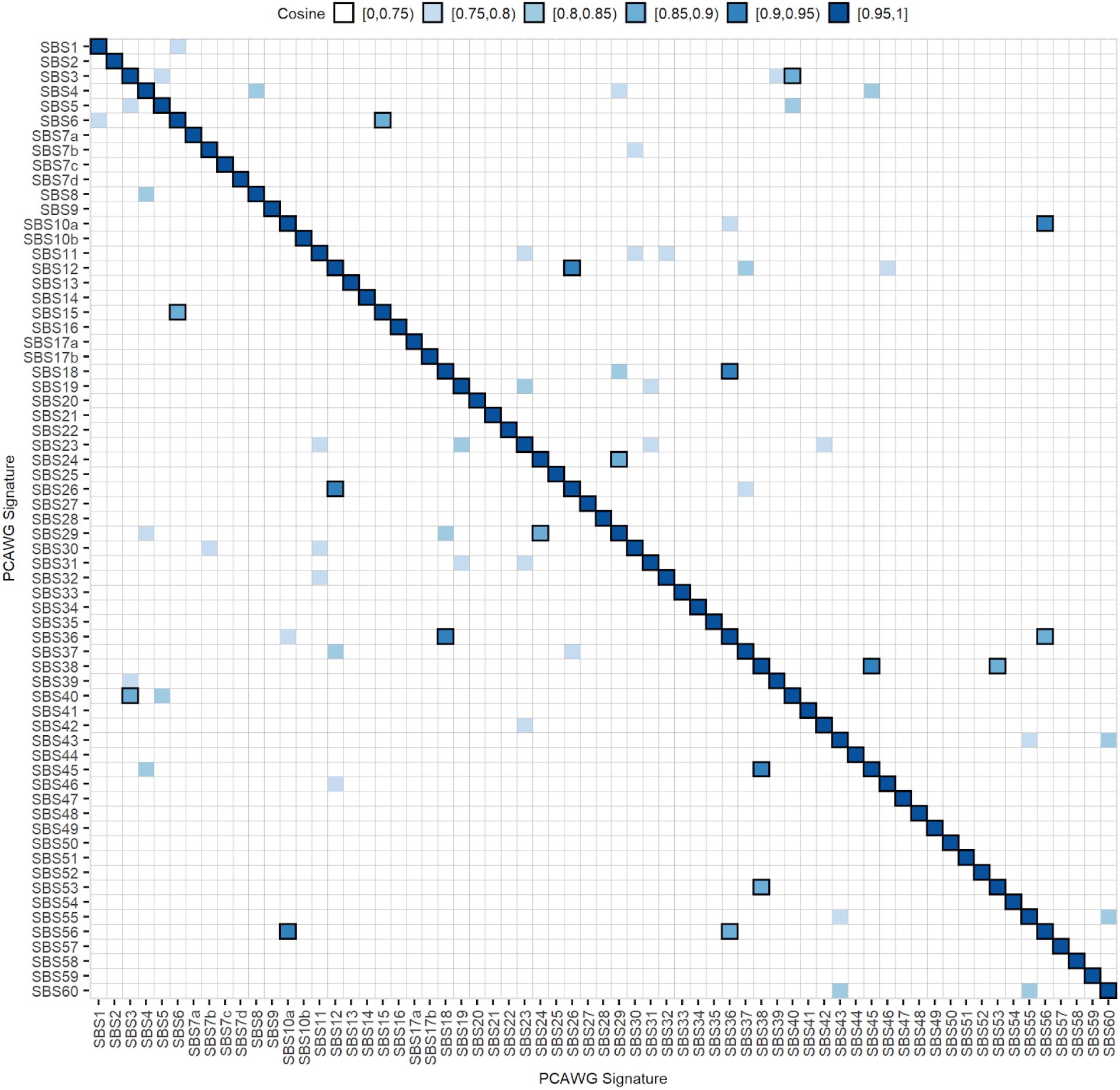
The known tumor mutational signatures may have similar mutation spectra to other known tumor mutational signatures. Cosine similarity between known tumor mutational signatures^12^ shows that some of the reported signatures are highly similar to each other, such as SBS6 and 15 (cosine=0.86), or SBS12 and 26 (cosine=0.93). Cosine similarities greater than 0.85 are marked with black borders.

**Supplementary Figure 6.**
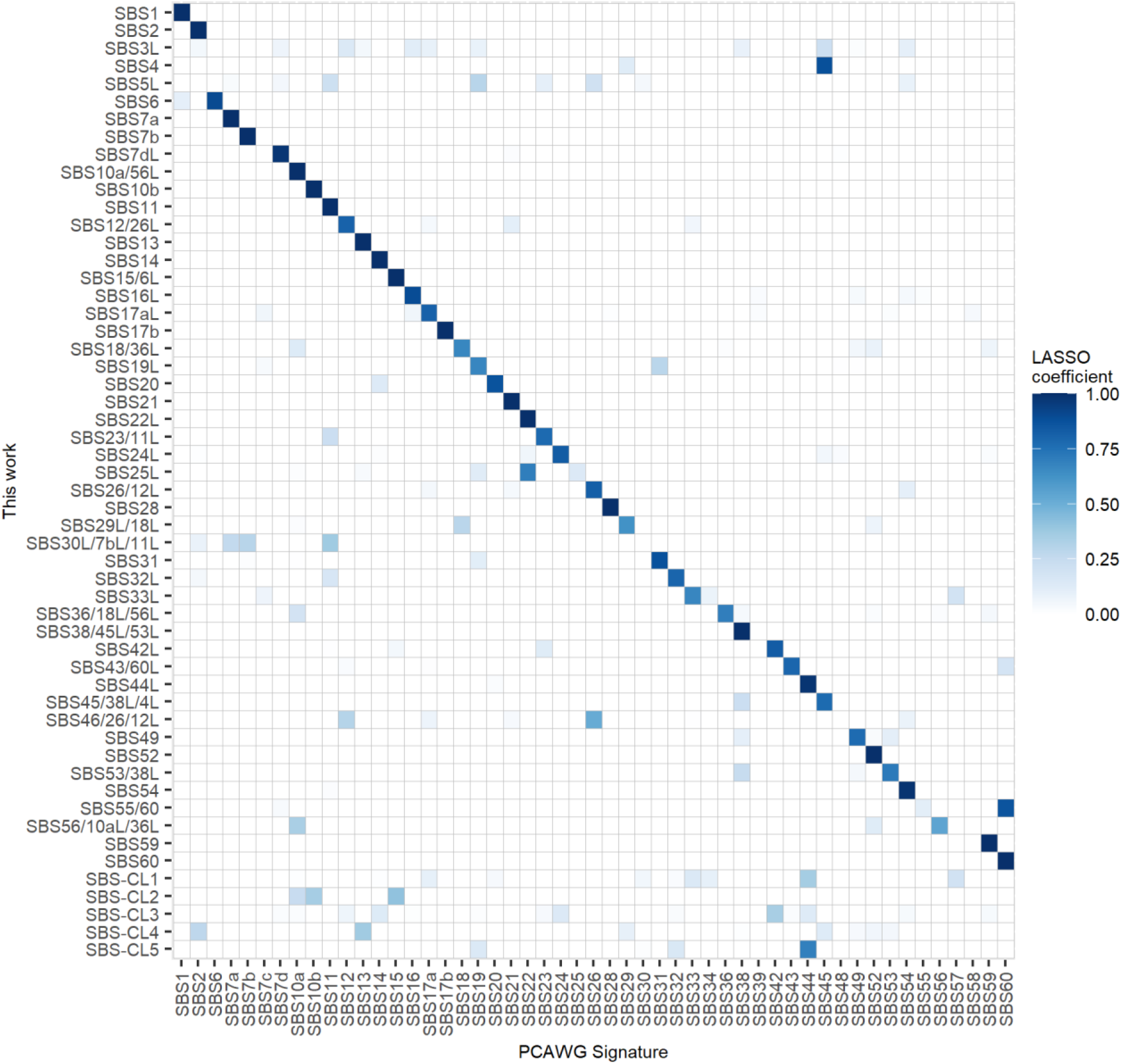
Modelling cell line mutation signatures as mixtures of tumor mutation signatures. Coefficients of LASSO linear regression where cell line mutational signatures are represented as a linear combination of known tumor mutational signatures reported by the PCAWG consortium^12^. For example, the 96-nucleotide spectrum of our cell line SBS-CL4 (penultimate row) may be modelled as a mixture of tumor SBSs (columns) 2, 13 and minor contributions of other signatures (primarily 29 and 45), while SBS-CL5 may be modelled as a mixture of SBS19, 32 and 44.

**Supplementary Figure 7.**
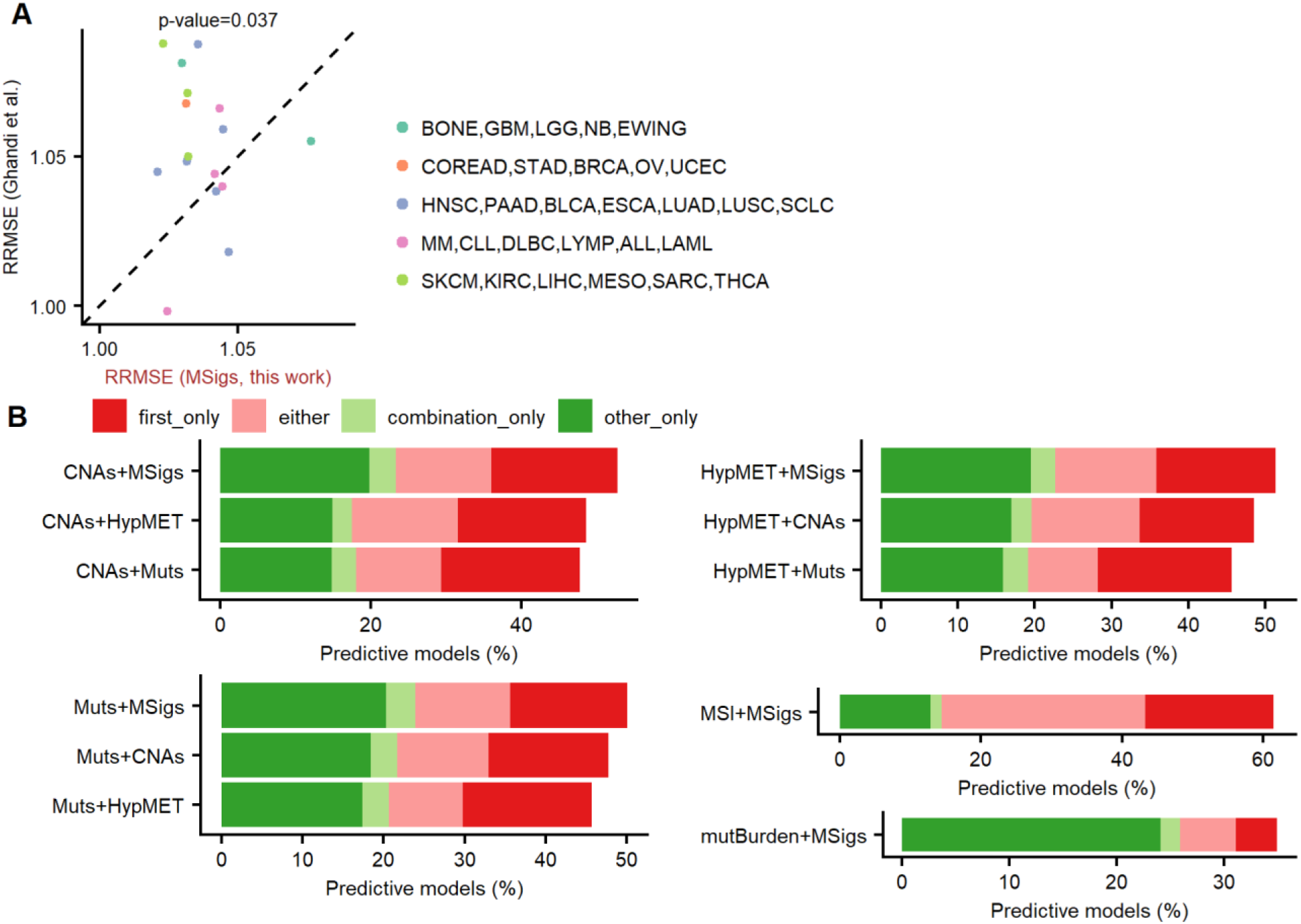
Complementarity of drug response prediction with mutational signatures and other molecular data types. (A) Predictive performance (RRMSE) of drug response prediction using Random Forest models based on mutational signatures (MSigs) reported here and previously^33^. The 667 overlapping cell lines from 16 cancer types between this study and the study of Ghandi *et al*. were considered. P-value of paired Wilcoxon signed rank test (one-sided) is reported. The dashed line denotes the diagonal. (B) Complementarity between mutational signatures reported here and other data types (copy number alterations, oncogenic mutations, DNA hypermethylation, the MSI/MSS status of cell lines, and a total mutational burden of cell lines. For each combination of data types, bars show the percentages of Random Forest models that are predictive (below-baseline RMSE) with mutational signatures but not another data type (“other_only”), by other feature type but not with mutational signatures (“first_only”), individually by either data type (“either”), or only by a combination of both data types (“combination_only”). The MSI+MSigs bar shows results only for 5 cancer types (colorectal, ovarian, stomach, uterine, and acute/chronic lymphoblastic leukemia) where MSI/MSS labels were available, while other panels show results for all cancer types.

**Supplementary Figure 8.**
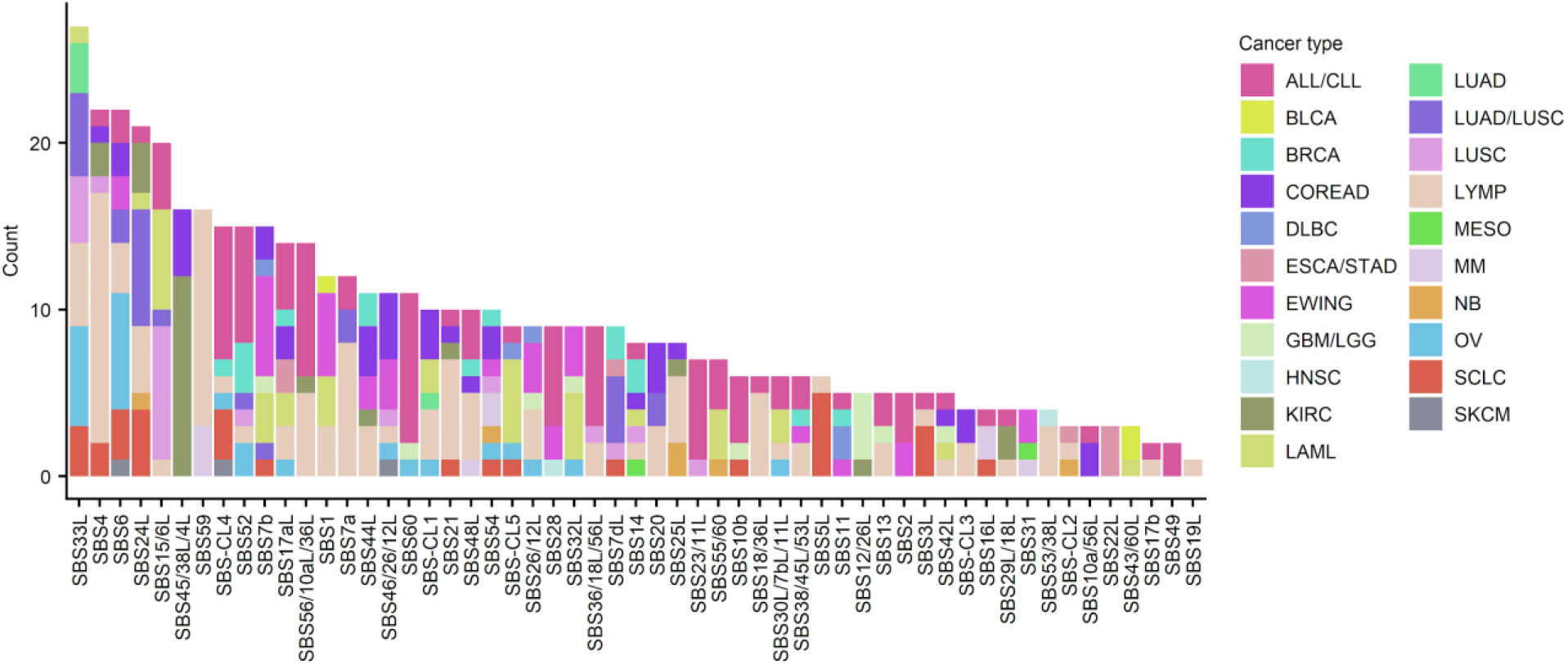
Overall tally of the statistically significant associations with continuous mutation signatures. Distribution of statistically significant associations (FDR<25%) across different cancer types.

**Supplementary Figure 9.**
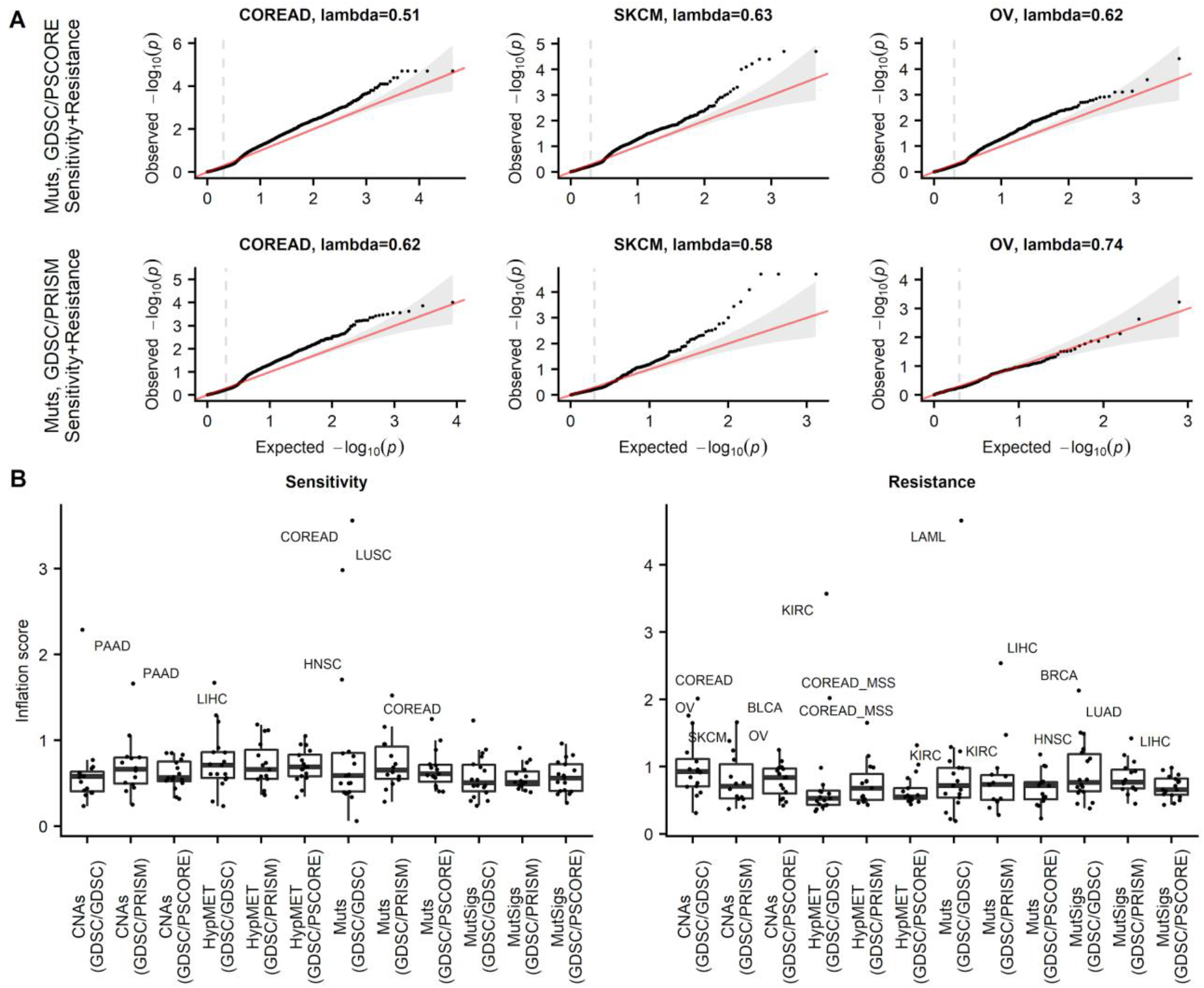
Calibration of p-values in the randomization tests for replication of drug-marker associations. (A) Examples of q-q plots for selected cancer types for GDSC/PSCORE and GDSC/PRISM tests. The inflation factor (lambda) is denoted for each qqplot. The vertical dashed line denotes the median. (B) Inflation factors across all cancer types, two-way tests and feature types (Muts, CNAs, HypMET, and mutational signatures) generally exhibit a deflation of p-values (lambda<1), while a few cancer types have inflated p-values (lambda>1.3; cancer types denoted on the plot) in some two-way tests and with some feature types.

**Supplementary Figure 10.**
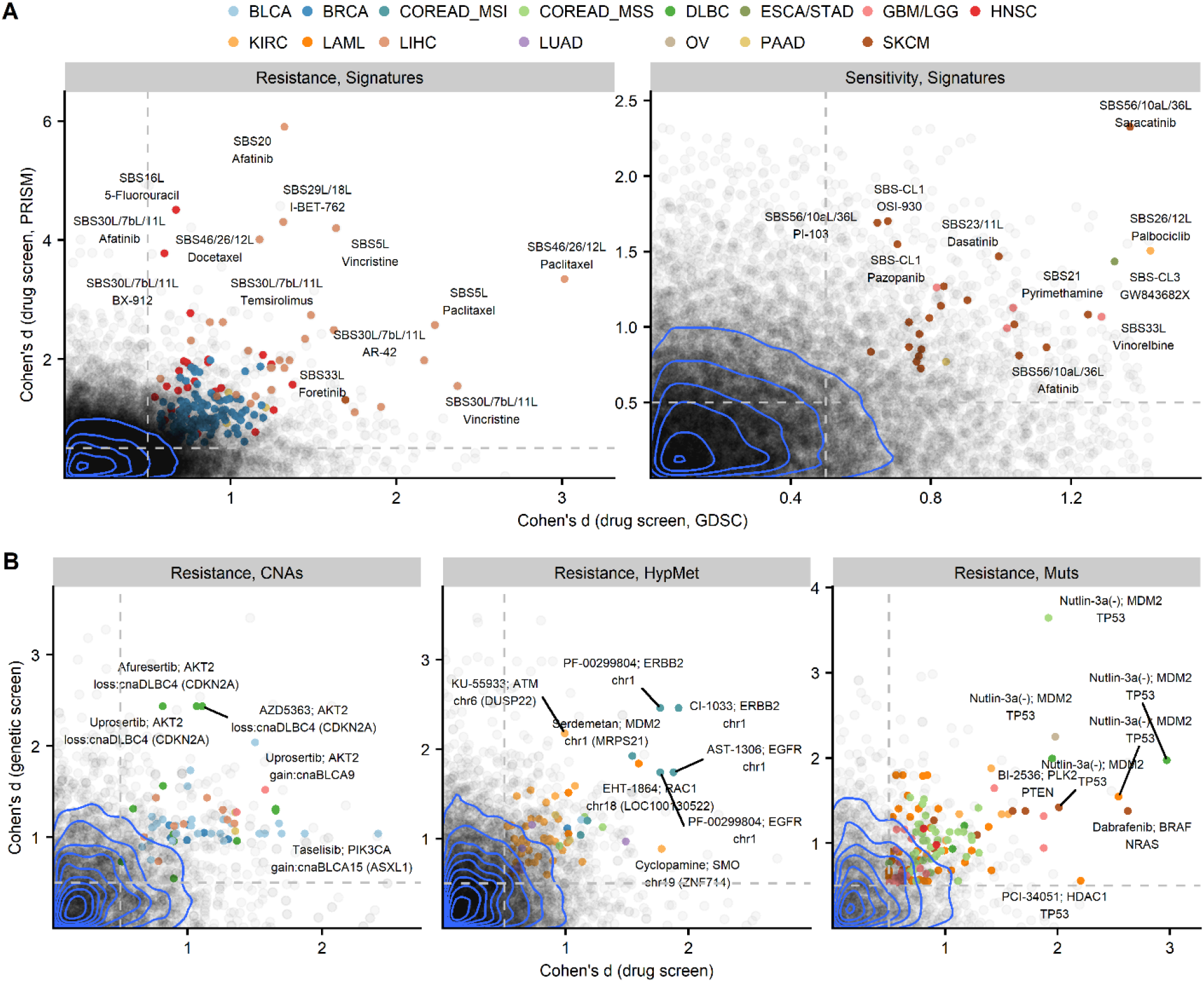
Associations that replicate in independent datasets. (A) The GDSC/PRISM replication test, which requires that the association between mutational signature and drug sensitivity or resistance replicates in the PRISM data set based on pooled cell line screening for drug response. Gray points represent all tested associations, while colored points denote the statistically significant associations according to a randomization test (Methods) and that also meet the required effect size threshold of Cohen’s d>0.5. Blue lines are the contours of the 2D kernel density estimates. (B) Plots as in A, but for replication of drug resistance associations (X-axis) in genetic screening data (Project SCORE, Y-axis) for classical markers: oncogenic mutations (Muts), copy number alterations (CNAs) and DNA hypermethylation (HypMet).

**Supplementary Figure 11.**
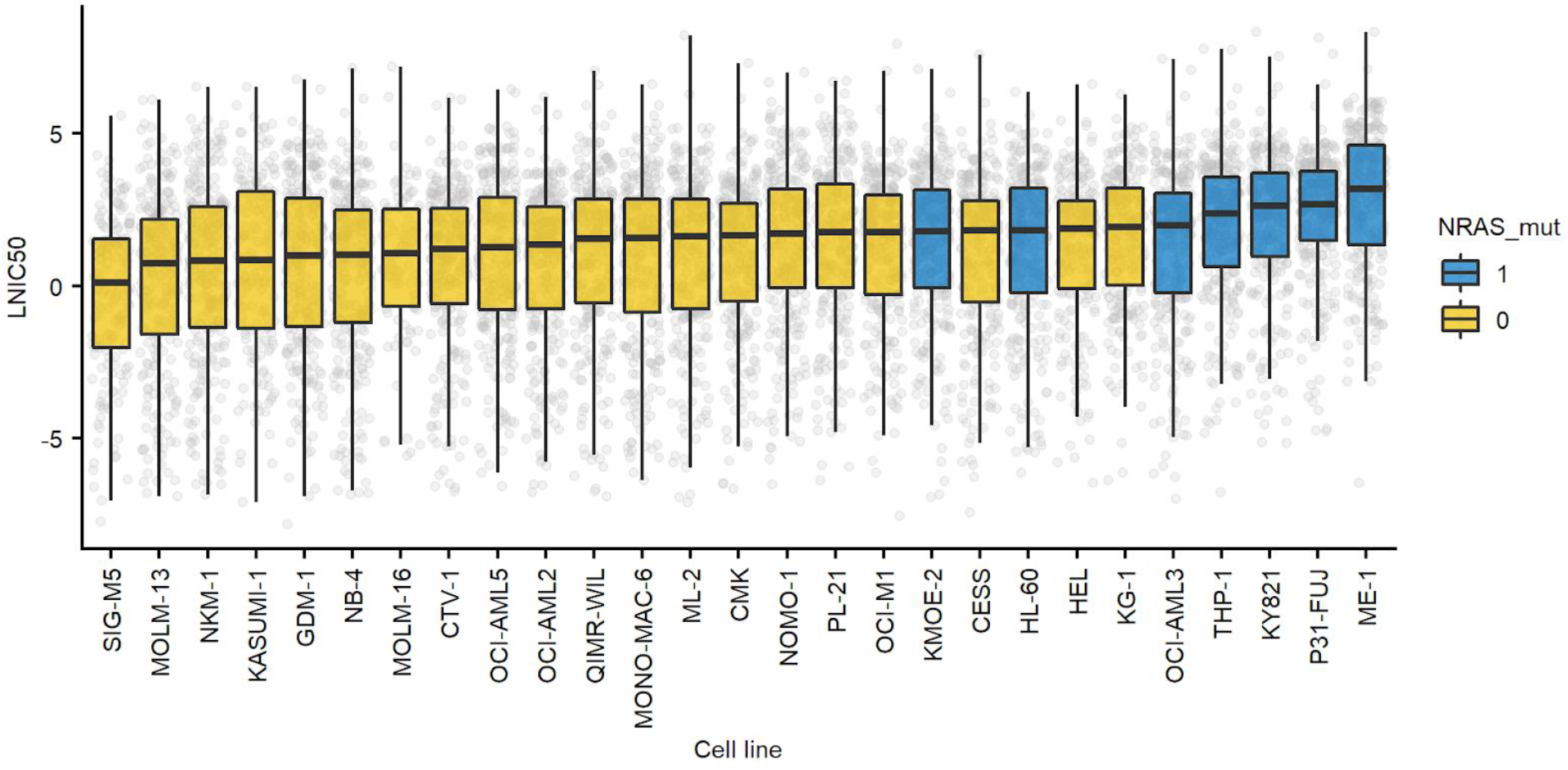
Sensitivity of acute myeloid leukemia cell lines to diverse drugs. Cell lines are sorted by median log IC50 calculated across all drugs screened against a cell line. NRAS mutant cell lines show (blue) increased median resistance to drugs compared to NRAS wild-type cells.

**Supplementary Figure 12.**
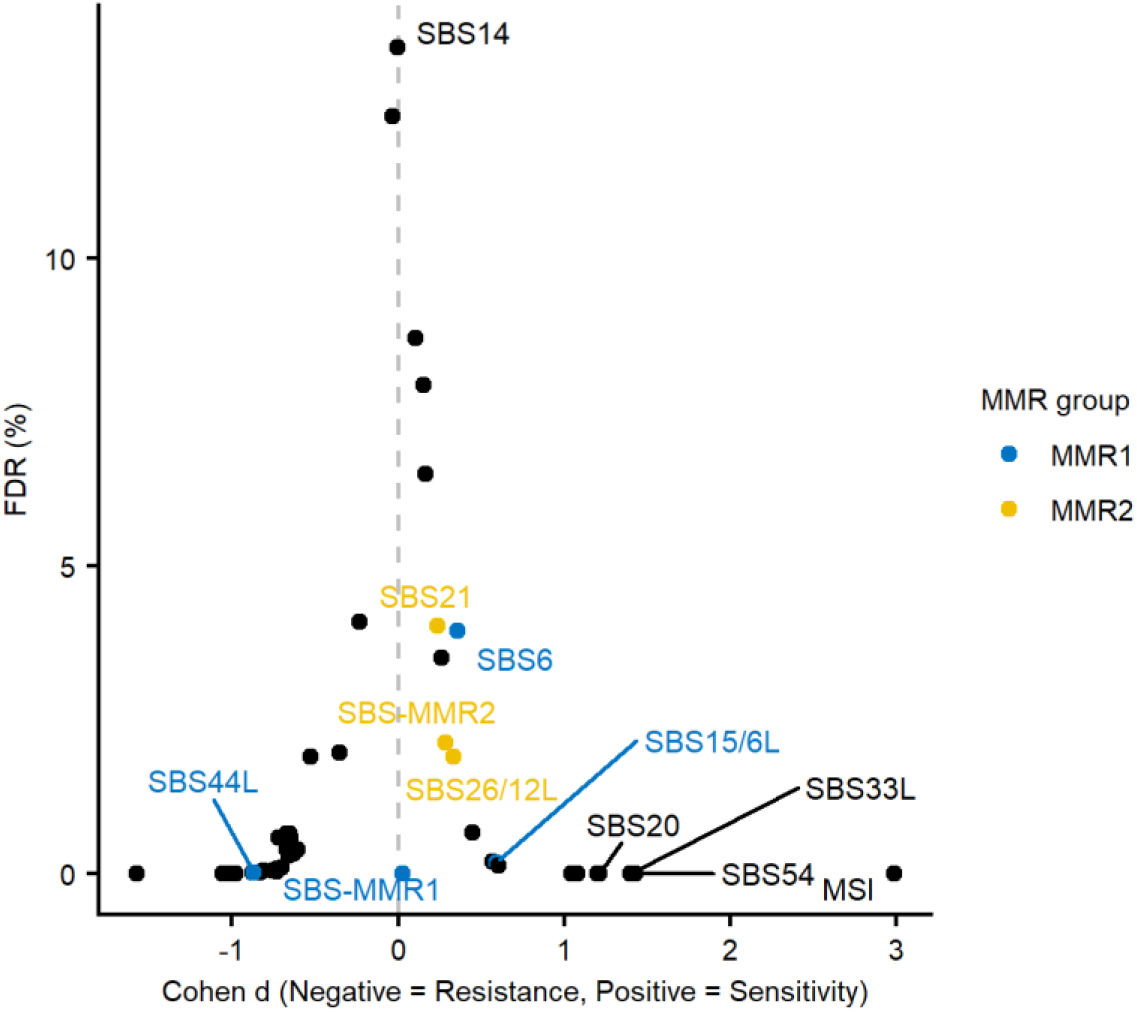
Different MMR-failure associated mutational signatures predict sensitivity to knock-out of the WRN gene to different extents. FDR of associations between different mutational signatures (including the MSI status of cell lines) and knock-out of the WRN gene in a pan-cancer analysis. The effect size is Cohen’s d statistic.

**Supplementary Figure 13.**
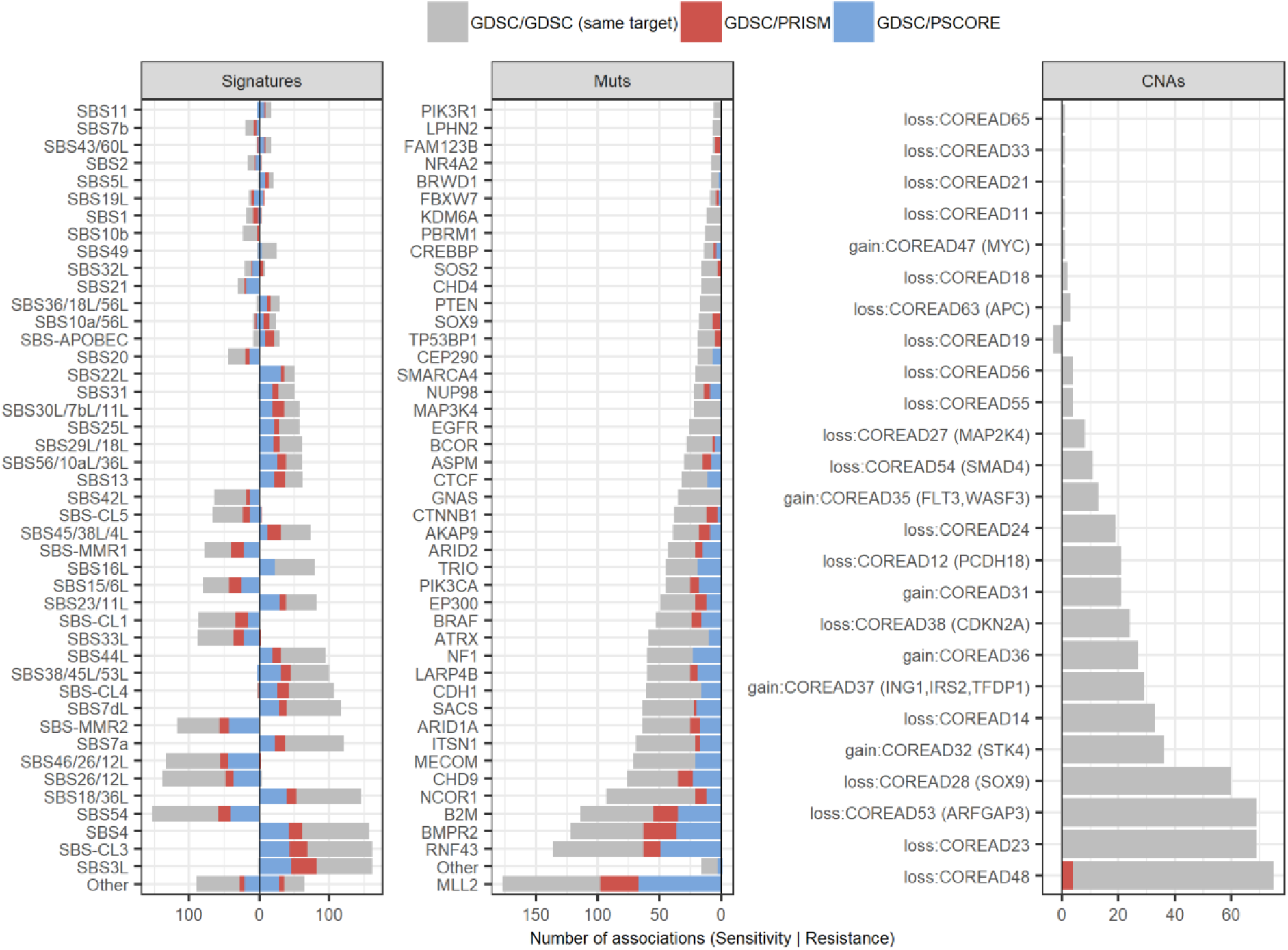
Tally of the significantly replicated associations with mutation signatures and other markers for colorectal cell lines without stratifying by MSI status. Comparison of the number of statistically significant associations (FDR<25%, effect size d>0.5) per feature, among mutational signatures, oncogenic mutations and copy number alterations in the three replication tests for colorectal cell lines (COREAD). Features are ranked by the number total of significant associations, either for drug sensitivity (negative side of X-axis) or resistance (positive side of X-axis).

**Supplementary Figure 14.**
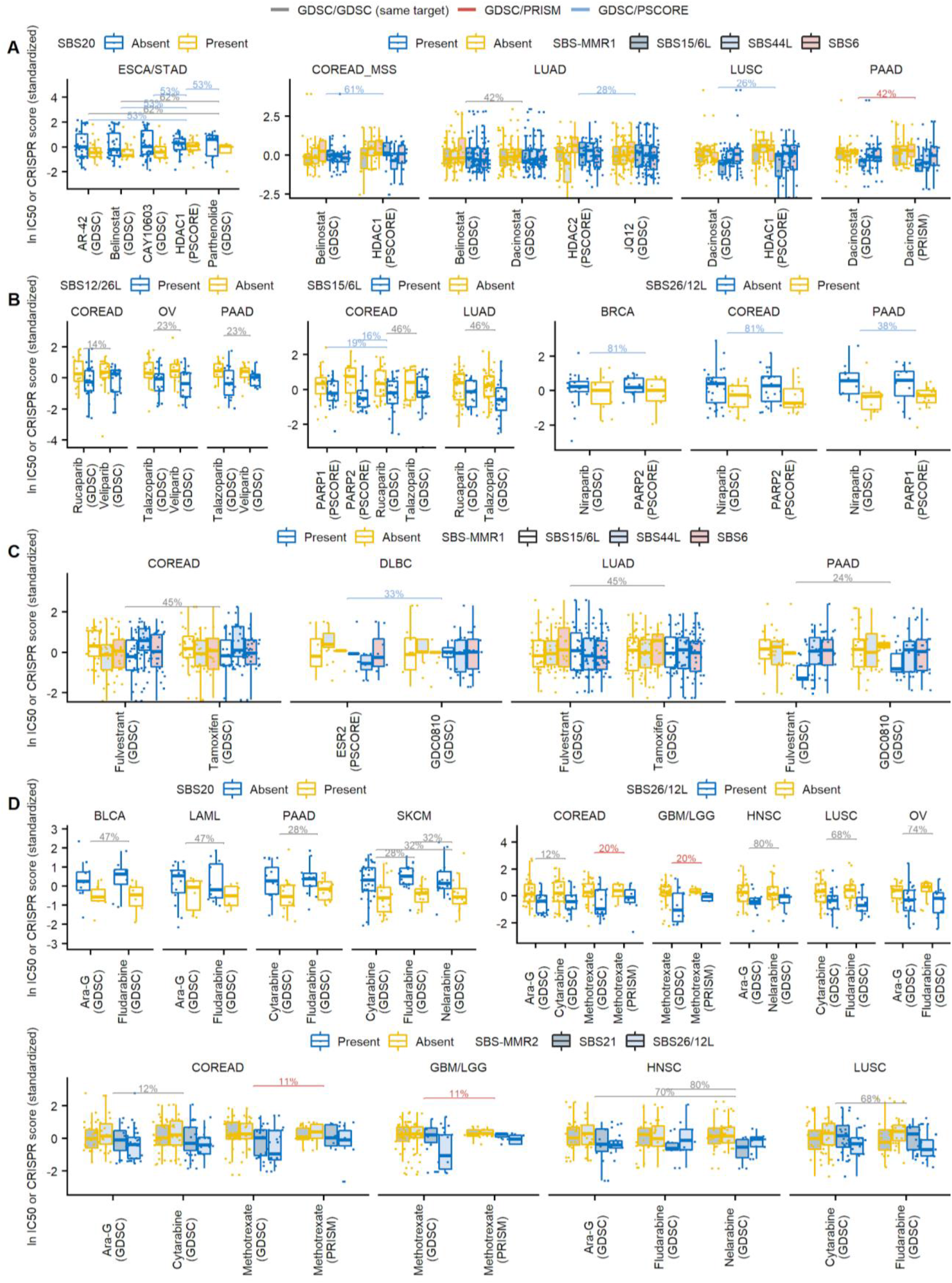
Additional examples of significant associations involving mutational signatures that replicate in independent datasets. (A) HDACi/SIRT example, accompanying the main Figure 5B. (B) Association of SBS20, 26/12L and MMR2 with sensitivity to DNA antimetabolites.

**Supplementary Figure 15.**
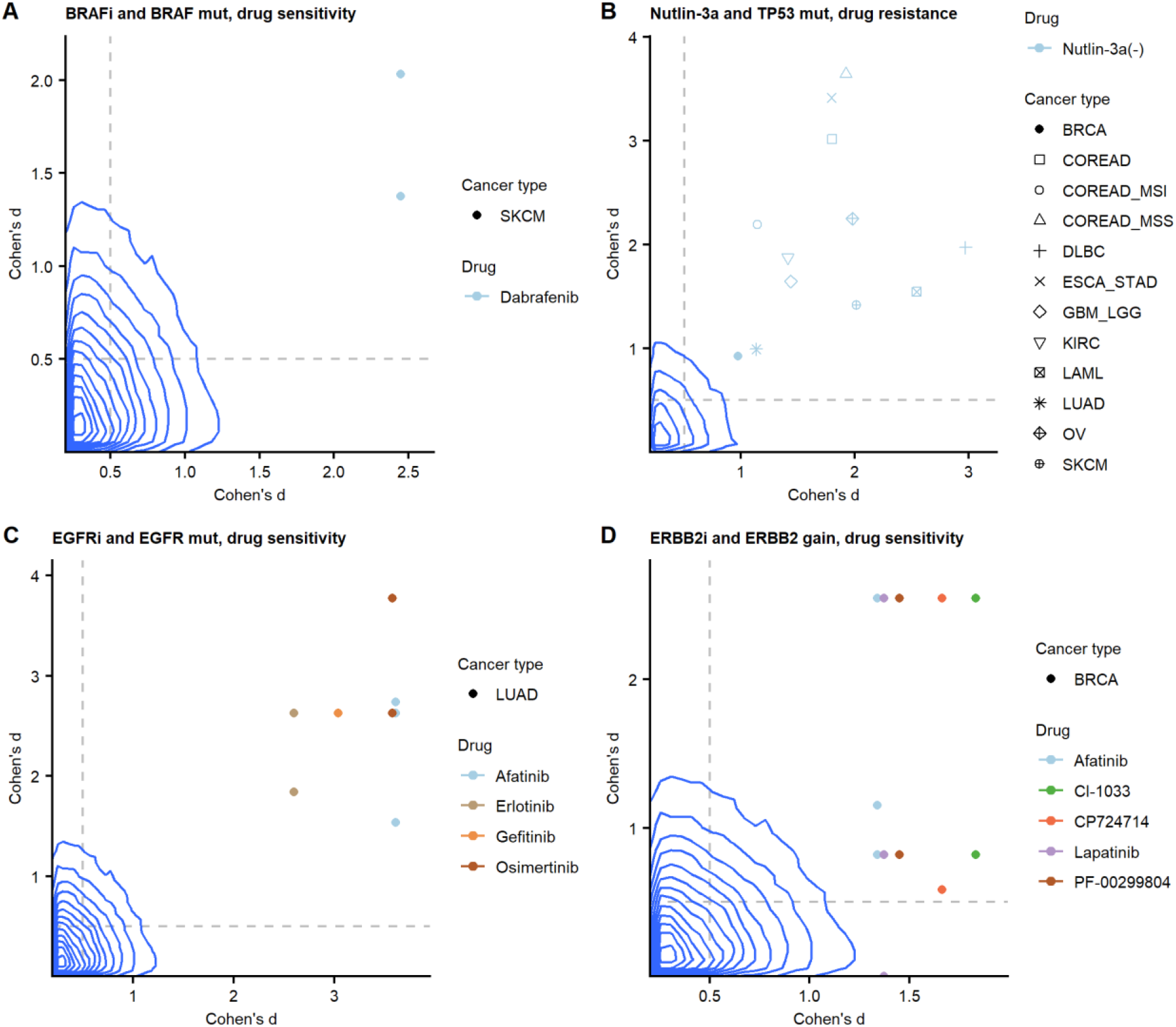
Drug sensitivity associations from ‘two-way’ replication tests involving known positive controls. (A) Sensitivity association between BRAF mutation and BRAF inhibitor Dabrafenib in melanoma cells. (B) Resistance associations between TP53 mutation and Nutlin-3a detected in several cancer types. (C) Sensitivity associations between EGFR mutation and EGFR inhibitors in colorectal and lung adenocarcinoma cells. (D) Sensitivity association between ERBB2 gain and several ERBB2 inhibitors in breast cancer cells. Blue lines are the contours of the 2D kernel density estimates of the distribution of associations between all drugs and mutations in all examined cancer genes (panels A, B, C) or all drugs and all recurrent copy number alterations (panel D). The dashed lines denote the 0.5 effect size threshold.

